# Motion processing in visual cortex of maculopathy patients

**DOI:** 10.1101/2025.02.07.636620

**Authors:** Célia Michaud, Jade Guénot, Cynthia Faurite, Mathilde Gallice, Christophe Chiquet, Nathalie Vayssière, Isabelle Berry, Yves Trotter, Vincent Soler, Carole Peyrin, Benoit R. Cottereau

**Affiliations:** CerCo UMR 5549, CNRS – Université Toulouse III, 31059 Toulouse, France; Univ. Grenoble Alpes, Univ. Savoie Mont Blanc, CNRS, LPNC, 38400 Saint-Martin d’Hères, France; Department of Ophthalmology, Grenoble Alpes University Hospital, 38700 La Tronche, France; IPAL, CNRS IRL 2955, 117418 Singapore

## Abstract

Previous studies on animal models suggested that visual areas involved in motion processing could undergo important cortical reorganizations following retinal damages. This could have major implications for patients suffering from macular degeneration (MD), one leading cause of vision loss. Here, we performed fMRI recordings in a group of maculopathy patients (including individuals suffering from age-related macular degeneration or from Stargardt’s Disease) and a control group to characterize the motion processing cortical network in MD patients and determine whether this network is modified following the onset of the scotoma. We used an experimental protocol based on random-dot kinematograms (RDKs) classically employed to characterize motion-selective areas in the brain. To ensure that the visual information processed by the two groups was equivalent, the visual field in each control participant was masked using an artificial scotoma directly derived from clinical measurements in their paired patient. We found that in MD patients, translational motion elicited significant and robust activations in a restricted cortical network which included the human V5/MT+ complex (hMT+), areas V3A and V6, and a portion of primary visual areas (V1, V2 and V3) connected to peripheral vision. Importantly, the same patterns of responses were also observed in control participants. Moreover, the extent and strength of activation within these motion-selective areas did not differ significantly between the two groups. Altogether, these results suggest that in humans, the motion-selective network does not undergo significant large-scale cortical reorganizations following the onset of MD.

**Significance statement:** Motion processing in the visual cortex of patients with macular degeneration has never been characterized. Here, we performed fMRI recordings in 7 maculopathy patients and found robust motion-selective activations in a cortical network which included the human V5/MT+ complex (hMT+), areas V3A and V6, and a portion of primary visual areas connected to peripheral vision. These activations closely align with those reported in participants with normal vision in the literature and do not significantly differ from those measured in a group of age and gender-matched control participants who viewed the motion stimuli with a matched artificial scotoma. Altogether, our results suggest that the motion-selective network does not undergo significant large-scale reorganizations in maculopathy patients following the onset of the scotoma.

## 1. Introduction

Macular degeneration (MD) is a visual pathology that alters the outer layers of the central retina, resulting in central vision loss. Its most common form is age-related macular degeneration (AMD) which develops after 60 years and currently affects over 67 million of patients in Europe (Li et al., 2020). Macular degeneration can also affect younger populations under the form of Stargardt disease (STG) which usually onsets during adolescence but can also appear later in life (Late-Onset Stargardt Disease; Westeneng-van Haaften et al., 2012). Despite their different origins, both diseases result in a loss of visual acuity and decreased contrast sensitivity, with severe consequences on patient’s quality of life as they impair recognition and mobility (Hassan et al., 2002; Kuyk & Elliott, 1999; Marron & Bailey, 1982; Wood et al., 2009). MD patients usually rely on peripheral vision, which has a much lower spatial resolution compared to central vision, to process their visual environment (Musel et al., 2011; Peyrin et al., 2017; Tran et al., 2011). Most of them develop a preferred retinal locus (PRL) outside the scotoma for ocular fixation (Fletcher & Schuchard, 1997). This overuse of peripheral vision was hypothesized to lead to cortical reorganizations in visual cortex (Baker, 2005), notably in neural populations which previously processed central vision. However, the associated neural mechanisms remain poorly understood. Most studies on cortical reorganisation in MD patients focused on the primary visual cortex (see however Little et al., 2008 or Ramanoël et al., 2018). Recordings using functional magnetic resonance imaging (fMRI) demonstrated that stimuli presented in peripheral vision activated portions of area V1 connected to the scotoma (Dilks et al., 2009), but it is now accepted that this effect is dependent on the cognitive tasks and low-level visual features, and only reflect unmasking of cortical feedbacks (Masuda et al., 2008, 2021; Morland, 2015). Interestingly, recent studies on animal models suggested that cortical reorganization after MD can be more pronounced in higher-level visual cortex, notably along the dorsal pathway. Using fMRI, Shao et al., (2013) found that visual responses in area MT, a dorsal region processing motion, were significantly modified in a macaque monkey with juvenile MD. In a cat model, Burnat et al., (2017) showed that the posteromedial lateral suprasylvian area PMLS (the homologue of primate cortical area V5/MT; Villeneuve et al., 2006) and to a lesser degree area 7 (which is also motion sensitive; Pigarev & Rodionova, 1998) had enhanced activity following the induction of a central retinal lesion in adult animals. These enhancements were associated with better performances in a motion task. In humans, a recent study showed that functional connectivity between the portions of early visual areas (V1, V2, and V3) that respond to peripheral vision and MT was enhanced in individuals with central vision loss (Fleming et al., 2024). Another study in psychophysics reported that self-induced motion perception from peripheral stimuli was increased in MD patients (Tarita-Nistor et al., 2008), suggesting some degree of plasticity in cortical areas processing motion. It is thus possible that neural responses in the motion-selective cortical network of MD patients are enhanced following the onset of their scotoma but this hypothesis has never been tested using functional neuroimaging.

Here, we employed an fMRI protocol commonly used to identify cortical areas selective to motion (see e.g., R. Tootell et al., 1995 or Zeki et al., 1991) in 7 MD patients and 7 age and gender-matched controls, whose central visual field was masked using simulated scotomas of the same positions and dimensions as those of their corresponding patients. Our main objectives were to characterize the motion processing cortical network in MD patients and to determine whether this network undergoes significant large-scale reorganisations following the onset of the scotoma. We tested if the cortical areas within this network are similar or different between the two populations and whether their associated BOLD responses are comparable. To complete our analysis, we performed psychophysical measurements to assess the motion discrimination thresholds in both groups.

## 2. Methods

### 2.1 Participants

Seven maculopathy patients (age range: 50 - 81) were recruited from the Retina Center of the Purpan Hospital of Toulouse and from the Ophthalmology center of the Grenoble Hospital (CHUGA) in France (see Table 1). Five of them were diagnosed with age-related macular degeneration (AMD), while the two others suffered from late-onset Stargardt disease. They all had dense central bilateral scotomas and a stable preferred retinal locus in at least one eye. None of them suffered from other visual illnesses. The experiments were also conducted in 7 age-and gender-matched controls with normal (or corrected to normal) vision. Age differences between patients and controls were all less than two years except for the control of MD2 (81) who was 76 (despite a broad screening procedure, we did not manage to find an older volunteer). Importantly, each control participant underwent the experiments in the same center as their paired patient. None of the participants suffered from general long-term illnesses. In order to evaluate their cognitive functions and their psychological states, they all underwent the Mini-Mental State Examination (MMSE) and the Beck Depression Inventory (BDI-II), respectively. All of them had a score above 23 for the MMSE and below 19 for the BDI-II. All participants gave their written informed consent and the protocol was approved by the French Ethics Committee (CPP) of Sud Mediterranée IV (2022-00957-36).

**Table 1:**
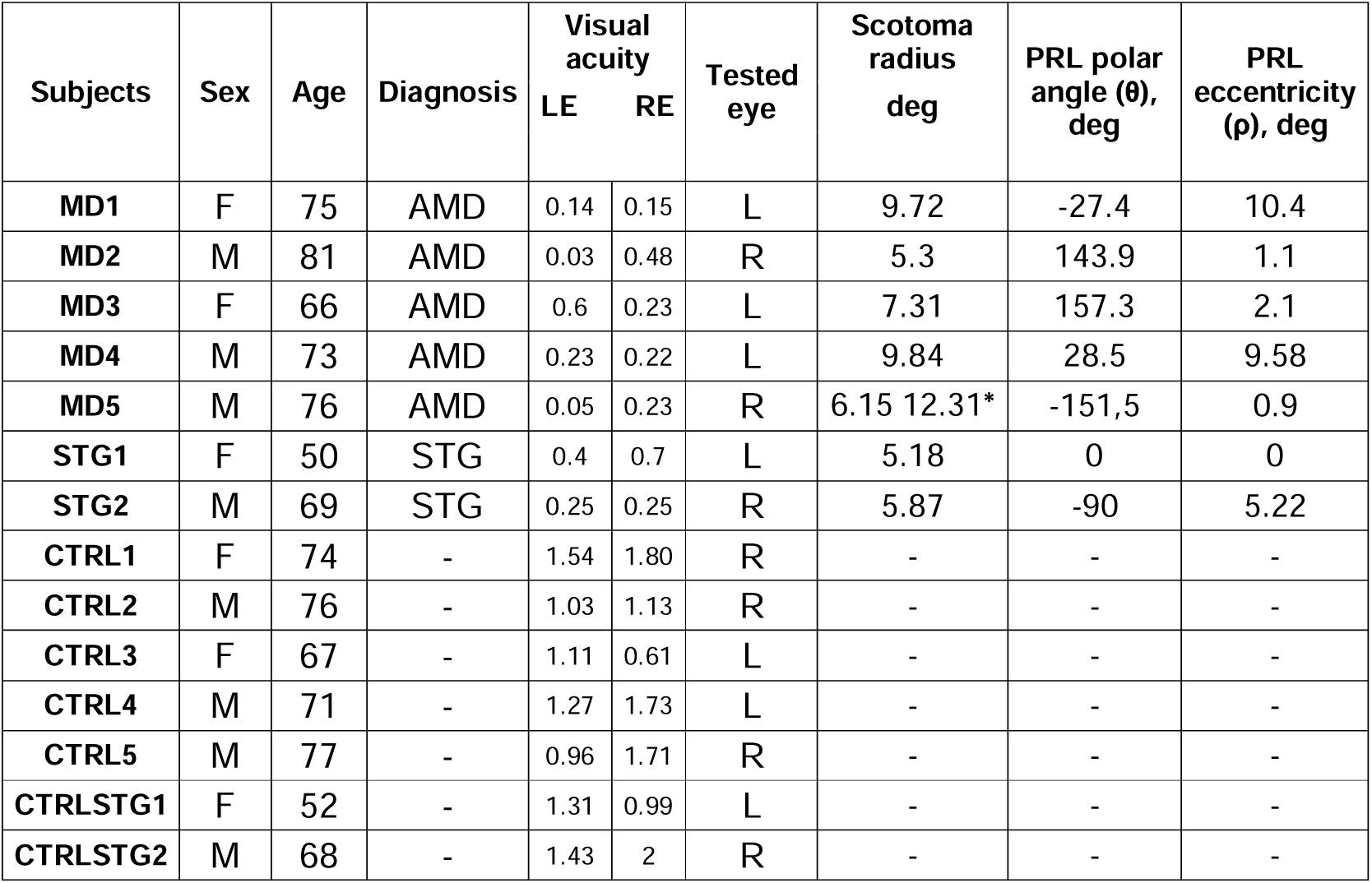
Characteristics of MD patients and age-matched controls included in the study. Precisions about the diagnosis of patients (AMD for Age-related Macular Degeneration, STG for late-onset Stargardt’s disease), the decimal visual acuity for the left (L) and right (R) eye, as well as the polar coordinates (angles and eccentricities) of the Preferred Retinal Locus (PRL) are included. Patients and their controls carried out the experiments in the same centers, i.e in Toulouse or in Grenoble. Controls for MD patients 1 to 5 are labelled CTRL1 to CTRL 5. Controls for STG patients STG1 and STG2 are labelled CTRLSTG1 and CTRLSTG2. It should be noted that because MD5 had an elongated scotoma, we used an ellipse rather than a disk for the artificial scotoma of his control. In this case, the two values provided in the ‘scotoma radius’ column (marked by an asterisk) provide the lengths of the ellipse semi-minor and semi-major axes.

| Subjects | Sex | Age | Diagnosis | Visual acuity | | Tested eye | Scotoma radius deg | PRL polar angle ($\theta$), deg | PRL eccentricity ($\rho$), deg |
| --- | --- | --- | --- | --- | --- | --- | --- | --- | --- |
|  |  |  |  | LE | RE |  |  |  |  |
| MD1 | F | 75 | AMD | 0.14 | 0.15 | L | 9.72 | -27.4 | 10.4 |
| MD2 | M | 81 | AMD | 0.03 | 0.48 | R | 5.3 | 143.9 | 1.1 |
| MD3 | F | 66 | AMD | 0.6 | 0.23 | L | 7.31 | 157.3 | 2.1 |
| MD4 | M | 73 | AMD | 0.23 | 0.22 | L | 9.84 | 28.5 | 9.58 |
| MD5 | M | 76 | AMD | 0.05 | 0.23 | R | 6.15 12.31* | -151,5 | 0.9 |
| STG1 | F | 50 | STG | 0.4 | 0.7 | L | 5.18 | 0 | 0 |
| STG2 | M | 69 | STG | 0.25 | 0.25 | R | 5.87 | -90 | 5.22 |
| CTRL1 | F | 74 | - | 1.54 | 1.80 | R | - | - | - |
| CTRL2 | M | 76 | - | 1.03 | 1.13 | R | - | - | - |
| CTRL3 | F | 67 | - | 1.11 | 0.61 | L | - | - | - |
| CTRL4 | M | 71 | - | 1.27 | 1.73 | L | - | - | - |
| CTRL5 | M | 77 | - | 0.96 | 1.71 | R | - | - | - |
| CTRLSTG1 | F | 52 | - | 1.31 | 0.99 | L | - | - | - |
| CTRLSTG2 | M | 68 | - | 1.43 | 2 | R | - | - | - |

For each participant, the eye with the better acuity was determined using the Sloan letters of the Freiburg Visual Acuity Test (Bach, 2007). In most participants, this eye was stimulated during the experiments and the other eye was patched. Only patients MD1 and STG1 were stimulated in the other eye because it was the only one with a stable preferred retinal locus (PRL, see below). In their case, the eye with the better acuity was patched.

### 2.2 Scotoma size

For each patient, an ophthalmologist delineated the border of the scotoma from the fundus of the eye captured by standard and autofluorescence imaging (see Figure 1-A for MD3, other examples are provided in Supplementary figure 1-1). The area of the scotoma (in square degrees of visual angles) was extracted using MATLAB (MathWorks, Natick, MA, USA). This area was used to compute the simulated scotoma of the age-and gender-matched controls (see below).

**Figure 1:**
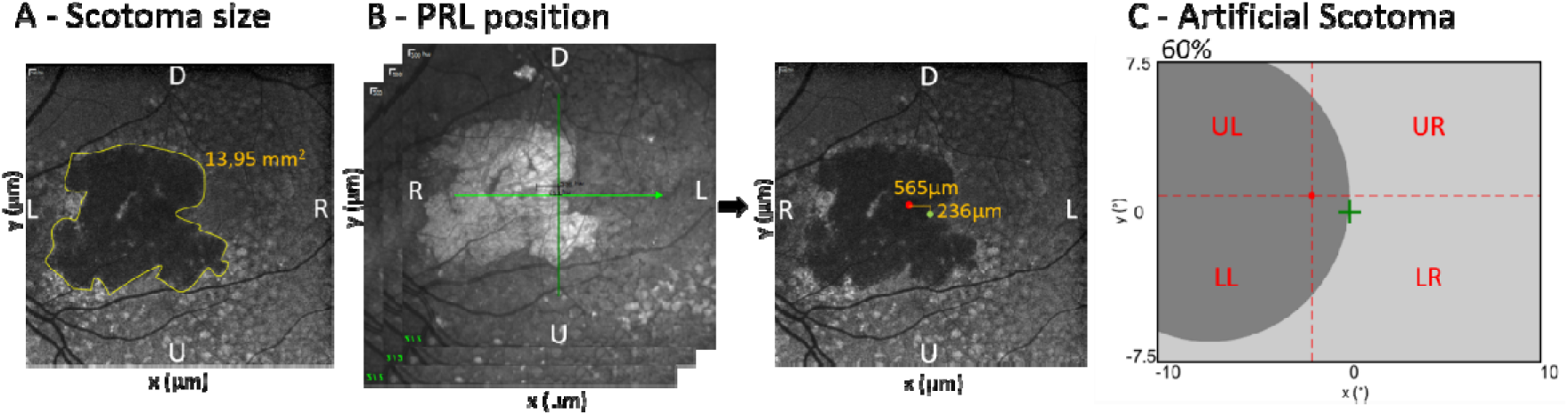
Estimation of the scotoma size and PRL position in one typical patient (MD3). These values are used to define the artificial scotoma masking the central visual field in the associated control participant. It should be noted that all retinal images in this figure have been inverted horizontally to be aligned with the visual field. The letters on panel A and B specify the orientation of the images (L: Left; R: Right; D: Down; U: Up). A) An autofluorescence imaging of the eye fundus is acquired. The border of the scotoma is delineated and the associated surface is used to calculate the radius of the scotoma in degrees of visual angle, given that 288µm on the image corresponds to 1 degree of visual angle. B) Patients were instructed to fixate a central light and we measured the x and y position of their PRL with spectral-domain optical coherence tomography (Spectralis Optical Coherence Tomography [OCT]; Heidelberg Engineering, Heidelberg, Germany). This process was repeated three times (leftward panel) and the final coordinates (green dot on the rightward panel) were the average across these three measures. Here, the position of the fovea is marked by a red dot. From the surface of the scotoma and the position of the PRL, it is possible to simulate the patient’s visual field. C) During the fMRI experiments, MD patients were instructed to fixate a central cross (shown here in green) with their PRL. Controls had to fixate a red cross at the position corresponding to their paired patient’s fovea while an artificial scotoma was displayed in dark gray. The dotted lines (not shown during the experiments) specify the four quadrants of the patients’ and controls’ visual fields in this case: upper-left (‘UL’), upper-right (‘UR’), lower-left (‘LL’) and (‘LR’). The screen (in light gray) dimensions provided here (20° x 15°) are those used in the two MRI centers. The percentage displayed on the top of the graphics specifies the visible portion of the screen (i.e., the surface that is not obscured by the artificial scotoma) in this case.

### 2.3 Position of the Preferred Retinal Locus (PRL) in maculopathy patients

MD patients usually develop a new locus of ocular fixation following the onset of their scotoma. To test for the presence of such preferred retinal locus (PRL) and to determine its location, we performed spectral-domain optical coherence tomography (see Figure 1-B). During the measure, patients were instructed to fixate a light produced by the OCT. Patients with a developed PRL naturally used it to direct their gaze, thereby allowing us to record its retinal coordinates. This process was repeated three times with a short break in between acquisitions, to ensure that the measure was stable (see Contemori et al., 2024). Table 1 reports the average PRL locations across these three measures (see Maniglia et al., 2018 or Guénot et al., 2022 for more details). It is important to note that two patients (MD5 and STG1) relied on central vision for fixation, indicating that they retained some residual visual function near their fovea despite significant damages to their maculas, as shown in Supplementary Figure 1-1. As will be demonstrated, the inclusion of these patients does not introduce biases into our fMRI results.

### 2.4 Artificial scotoma and viewing conditions in control participants

For each control participant, we masked the central visual field using an artificial scotoma as in previous studies (Christen & Abegg, 2017; Gupta et al., 2018). This simulated scotoma had the same surface as the one measured in the associated patient (see the ‘*Scotoma size*’ section) and was centered on the fovea (except for CTRL2 and CTRL 3 where the artificial was shifted to the left because MD2’s and MD3’s scotoma occupied a larger portion of the left visual hemifield). Control participants were instructed to gaze on a fixation cross that we displayed at the position of their paired patient’s fovea (see the red cross on Figure 1-C). In this case, the center of the screen fell on the visible part of their visual field, but at the PRL location of their paired patient. For all control participants, artificial scotomas were simplified as a disk except for CTRL5 for which we used an ellipse to more closely match the real shape of MD5 scotoma, which was very elongated (see Supplementary figure 1-1). This protocol permits patients and controls to receive comparable visual inputs. It notably equalizes the position and surface of the occluded part of the visual field (see Guénot et al., 2022).

### 2.5 Motion stimuli

Motion stimuli were adapted from the experimental protocol of a previous electrophysiological study (Graziano et al., 1994). They consisted in translational patterns defined from randomdot kinematograms (RDKs) (see Figure 2-A) generated with Matlab (R2017a). The RDKs contained bright non-overlapping dots (diameter: 0.2°) moving on a homogenous dark background with a high contrast (100%). They had a density of 0.3945 dots per square degree of visual angle. When a dot reached the border of the display screen, it was immediately relocated at a random position. This process permits to equalize mean luminance and dot density across the screen during the experiments. Each dot moved at a velocity of 7°/s, which corresponds to the average preferred speed of neurons in macaque MT (Priebe et al., 2006). This speed is in agreement with those used in previous fMRI studies on motion processing (6°/s in Zeki et al., 1991, 8°/s in (Huk et al., 2002). These translational motion stimuli were used for the fMRI recordings (see the next section) and for the psychophysical measurements (see the ‘*Experimental protocol for the psychophysical measurements*’ section).

**Figure 2:**
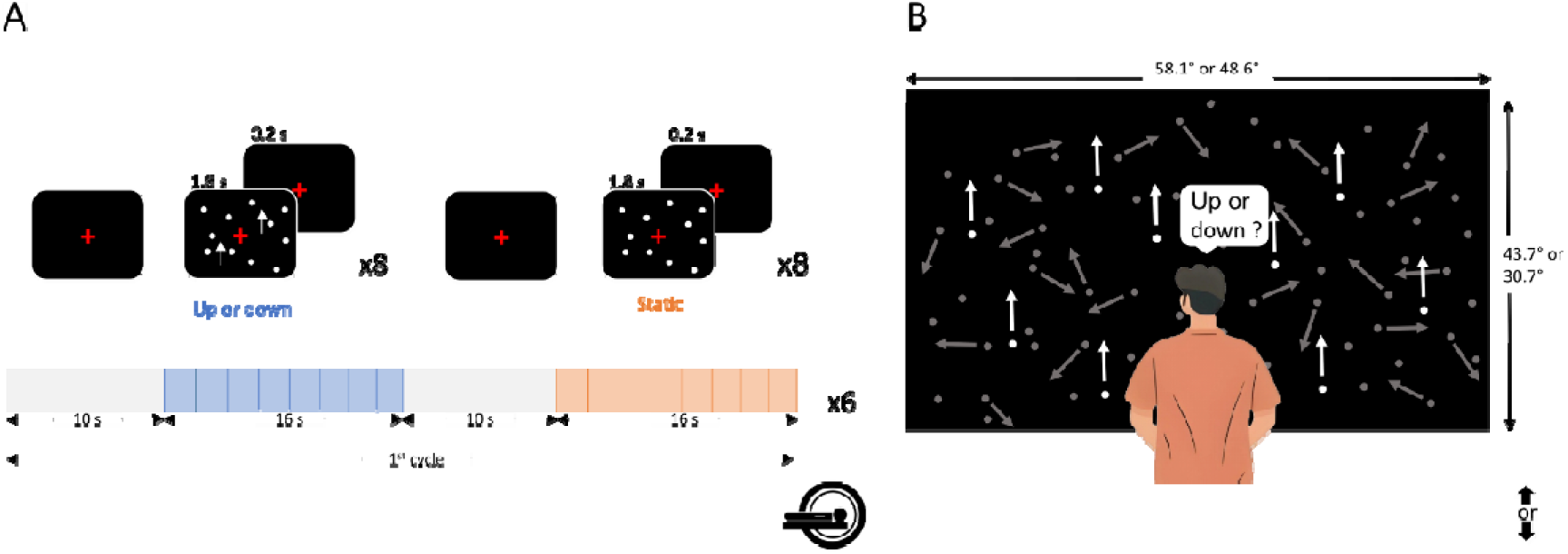
Experimental protocols. A) fMRI recordings were performed using a block-design. Runs lasted 312 s and contained 6 identical cycles of 52 s. Each cycle started with a baseline period of 10 s during which only a red fixation cross was displayed on a black screen. It was followed by a block of 16 s of uniform motion (shown here in blue), then by another 10 s of baseline and finally by a block of 16 s with static dots (in red). During motion blocks, a new pattern of translational motion during which all the dots moved either upward or downward was presented every 2 s. These patterns last 1.8 s and were separated by short periods (0.2 s) during which the screen went black and only the fixation cross was presented. The same principle was applied during the blocks of static dots (a new pattern was displayed for 1.8 s every 2 s and patterns were separated by 0.2 s). B) During psychophysical recordings, participants had to report the perceived motion direction of the stimuli (either upward or downward). We manipulated motion coherence and estimated thresholds corresponding to 80 percent of correct detections. For illustration, an upward motion trial is shown at 20% of coherence. Signal dots are shown in white and noise dots in gray for better visibility on the figure (all the dots were white during the experiment). During all the recordings (fMRI and psychophysics), participants had to fixate a red cross that was central for patients (who used their PRL) and at the position corresponding to their paired patient’s fovea for controls. This red fixation cross was always present on the screen.

### 2.6 Experimental protocol for fMRI measurements

In both Toulouse and Grenoble MRI platforms, stimuli were displayed on a large screen and covered 20°x15° of visual angle. fMRI recordings were performed using a block-design. Each run consisted of 312 s and were divided into 6 identical cycles of 52 s (see Figure 2-A). Each cycle started with a baseline period of 10 s during which only a red fixation cross of 1° of visual angle was displayed on a black screen. It was followed by a block of 16s of uniform motion, then by another 10 s of baseline and finally by a block of 16 s with static dots. During motion blocks, a new pattern of translational motion during which all the dots moved either upward or downward was presented every 2 s. These patterns last 1.8 s and were separated by short periods (0.2 s) during which the screen went black and only the fixation cross was presented. The same principle was applied during the blocks of static dots (a new pattern was displayed for 1.8 s every 2 s). During both translational motion and static dot blocks, an artificial scotoma masked the central vision of control participants.

This protocol is classically employed to identify motion selective cortical areas (Zeki et al., 1991; Watson et al., 1993; Tootell et al., 1995a; Huk et al., 2002). The visual display of the stimuli and its temporal synchronization with the acquisition of the fMRI data was controlled using the EventIDE software (Okazolab). A red fixation cross was displayed during all the functional recordings, in the center of the screen for patients (who directed their gaze with their PRLs) and at the position corresponding to their paired patient’s fovea for controls (see above). As shown in the central columns of Figures 3 and 4, the visible portion of the stimuli covered most of the screen for the majority of participants. The exceptions were MD5 and his control, for whom 42% of the screen was visible. In order to make sure that all participants correctly directed their gaze and were focused on the visual stimuli, a white circle (0.75° of radius) was randomly displayed for 200 ms on the center of the screen during the runs (10 displays per run on average). Its size and duration were determined from pilot experiments. Participants were instructed to press a button on a gamepad whenever they detected it. Even though this task was not easy (notably for patients who were fixating with their PRLs), they all performed it well (mean percentage of correct detection for patients: 80; and controls: 90) and we thus kept all the runs for further analyses.

**Figure 3:**
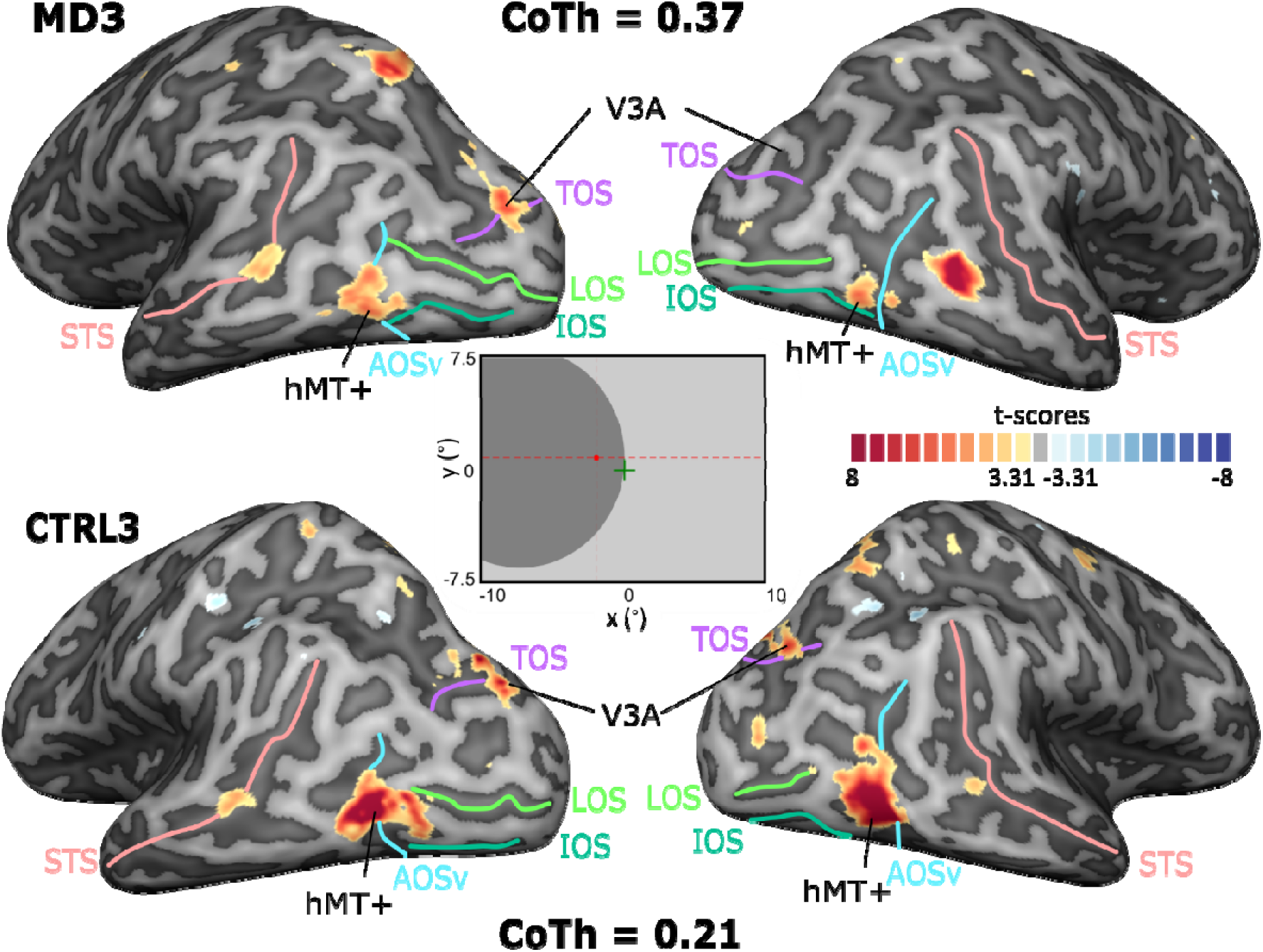
Comparison between BOLD responses to the translational motion condition and to its control condition (static dots) in one representative patient (MD3) and her age-and gender-matched control (CTRL3). Activations are shown on inflated individual cortical surfaces. Gyri appear in pale gray while sulci are shown in darker gray. Cortical nodes with significantly stronger responses to translational motion appear in yellow/red colors while nodes with significantly stronger responses to static dots are shown in blue. Data were thresholded at p-value < 10^-3^ (uncorrected). It should be noted that area V3A in the right hemisphere of MD3 was detected using a threshold p-value of 10^-2^ and it is thus not visible here. Colored lines provide the anatomical positions of the Inferior Occipital Sulcus (IOS, in olive green), of the Lateral Occipital Sulcus (LOS, in green), of the Transverse Occipital Sulcus (TOS, in purple), of the ventral part of the Anterior Occipital Sulcus (AOSv, in cyan), and of the Superior Temporal Sulcus (STS, in red). The central panel illustrates visual stimulation in this case. During the fMRI recordings, MD3 fixed the center of the screen (green cross) with her PRL while CTRL3 gazed at the position of MD3’s fovea (red cross). The dark disk is the artificial scotoma (see Figure 1 for more details on its definition). The dotted lines (not shown during the recordings) specify the positions of the horizontal and vertical meridians. We also provide the coherence thresholds (CoTh) estimated through psychophysical measurements.

**Figure 4:**
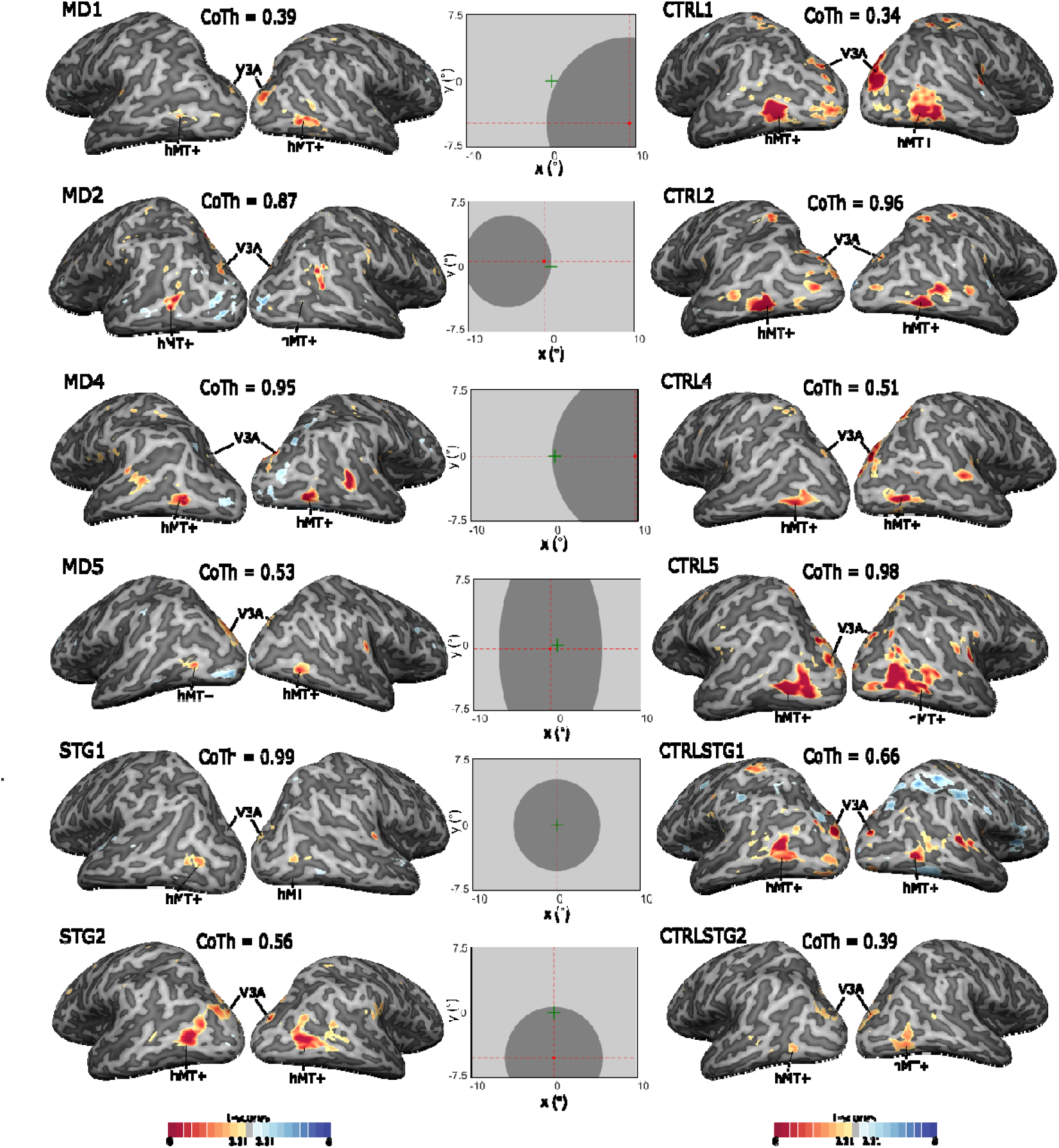
Comparison between BOLD responses to the translational motion condition and to its control condition (static dots) in all participants, except MD3 and her control (CTRL3), whose data are provided on enlarged views in Figure 3. Significantly stronger responses to motion are shown on lateral views of the cortical surfaces for MD patients (leftward panels) and their controls (rightward panels). See Figure 3 for the details of the legend.

### 2.7 (f)MRI recordings

(f)MRI recordings were performed at 3T (Achieva dStream 3.0T TX, Philips, NL) at Toulouse NeuroImaging Center (ToNIC, Toulouse, France) and at IRMaGe MRI facility (Grenoble, France). Importantly, both MRI platforms are equipped with the same MR hardware configuration (with Quasar Dual gradient system and dStream architecture supporting MultiBand-accelerated acquisitions, 32 channel head coil). They were operated using the same software version (R5.7.1). This permits to minimize the differences between the recordings performed in the two cities, although we emphasize that each patient from one center was matched with a control from the same center. We acquired two high-resolution anatomical images of the participant’s brain using a T1-weighted magnetization-prepared rapid gradient-echo (MPRAGE) sequence (repetition time, TR = 8.1 ms; echo time, TE = 3.7 ms, flip angle = 8°; FOV: 240 x 240 mm; voxel size = 1 x 1 x 1 mm; 170 sagittal slices). These anatomical images were first co-registered and then averaged together to be used as a reference to which the functional images from all the experiments were aligned. Functional images were collected with a T2 * weighted gradient-echo echoplanar imaging (EPI) sequence (TR = 2000 ms, TE = 30 ms, flip angle = 75°, SENSE factor = 1.6; multi-band (MB) = 3; FOV: 224 x 224 mm; matrix size: 112 x 110 mm; voxel size = 2 x 2 x 2 mm, 63 transverse slices, dyn = 156).

### 2.8 (f)MRI data analyses

MRIQC v.13.1.0 was used to check the quality of all our functional recordings (Esteban et al., 2017). We did not observe any out-of-range runs in our data, indicating negligible head-movement. We thus kept all the measurements for further analyses. Pre-processing of fMRI data included slice scan time correction, 3D motion correction using trilinear/sinc interpolation, and a linear trend followed by a high-pass filtering using Fourier analysis with a cut-off at 3 cycles. For each participant, functional data were co-registered on the anatomy. Functional and anatomical data were brought into anterior commissure - posterior commissure (ACPC) space using cubic spline interpolation and then transformed into standard Talairach (TAL) space (Talairach & Tournoux, 1988). For further analyses, the volume time courses were spatially smoothed at 4 mm FWHM.

Individual analyses were performed on each node of the participant’ cortical tessellations by fitting a general linear model (GLM) to the corresponding blood oxygen level-dependent (BOLD) signal. The model contained 2 main regressors, representing the 2 experimental conditions: uniform motion (either upward or downward) and control (static dots). These regressors were convolved with a model of the human hemodynamic response function. We also introduced 8 regressors of non-interest in our model. 6 of them corresponded to the translation and rotation of the head estimated during the motion-correction step of the pre-processing (see above). The two last ones were time-courses captured within the ventricles and the white-matter. This operation permits to reduce the noise in the data (see e.g., Audurier et al., 2022). For each participant, the beta weights obtained from the GLM were subsequently used to perform univariate analyses (t-scores) at the whole-brain level. All these analyses were realized using the BrainVoyager software (v23, Brain Innovation, Maastricht, the Netherlands) (Goebel et al., 2006). Results are shown on inflated surfaces, obtained by segmenting the white and gray matter, as well as the cerebrospinal fluid, and reconstructing a surface at the boundary of the white and gray matter. Statistical maps are mainly thresholded at p-value < 10^-3^ (uncorrected), as is often the case in individual-level fMRI analysis (see e.g., Aedo-Jury et al., 2020).

Group-level analyses were performed within independently defined regions of interest (ROIs). These ROIs were given by spheres in the volumetric space centered on the Talairach coordinates provided in previous studies for areas hMT+ (Watson et al., 1993), V3A (R. B. H. Tootell et al., 1997) and V6 (Pitzalis et al., 2010). Within each ROI, we computed the differences between the averages percentages of signal changes (PSCs) during the translational motion condition and during its control condition (static dots). For each group, we first determined whether these differences of PSCs were significantly different from zero across participants. We used Wilcoxon sign-rank tests because it is non-parametric and thus does not assume that the data are normally distributed. Then, we tested whether the differences of PSCS were different between the two groups (MD patients and controls), also using Wilcoxon sign-rank tests. Here, data were paired between each patient and his/her control to reflect the fact that each pair received the same visual stimulations. To complete our statistical analyses and determine how much evidence favors the alternate hypothesis (H_1_: the differences of PSCs are not the same between the two groups) against the null hypothesis (H_0_) in a given ROI, we also computed the Bayes factor using the R-code open source JASP program. This factor is given by the ratio of the likelihood of the alternate hypothesis (H_1_) to the likelihood of the null hypothesis (H_0_). Values weaker than one favors the null hypothesis whereas values greater than one favors its alternative (Raftery, 1995). We used spheres with a radius of 10 mm, a value which leads to a cortical surface in line with the one reported for area hMT+ (see Kolster et al., 2010). It should be noted that the analyses were also repeated with radius values of 5 and 15 mm to make sure that the choice of this parameter did not influence our results.

### 2.9 Experimental protocol for psychophysical measurements

The psychophysical measurements were performed separately from the fMRI recordings, and outside the MRI scanner. During the experiment, participants sat in a chair whose height was adapted in order to adjust the height of the eyes to the center of the screen. Their head was placed on a head-support device clamped on top of a table and equipped with both chin and forehead supports. The chair and head-support devices were positioned so as to ensure a fine alignment between the participants’ head and trunk axes. Participants had to keep this position as constant as possible. Dot stimuli of 0.2° in size were presented on computer screens using a large portion of the visual field (58.1° × 43.8° of visual angle in Toulouse and 48.6° x 30.7° in Grenoble) to maximize coverage of peripheral vision. As for the fMRI recordings, during the whole experiment, participants had to fixate monocularly with the same study eye (the other eye was patched) a red cross. This cross was central for patients (who used their PRL) and at the position corresponding to their patient fovea for age-matched controls.

We used the same random dot kinematogram stimuli as for the fMRI experiments (see the ‘*Motion stimuli*’ section), except that here, each dot had a limited lifetime of 200 ms (12 frames at 60 Hz). At the end of its lifetime, it was randomly reassigned to a new spatial position. To avoid a coherent flickering of the stimulus every 200 ms, each dot initial age was randomly picked between 0 and 166 ms (11 frames) at the beginning of each trial. We chose to manipulate lifetime to permit a direct comparison of the psychophysical results with those obtained in a previous study using the same paradigm, but performed on other MD patients (Guénot et al., 2022). It should be noted that we used unlimited lifetimes for the fMRI recordings in order to maximize the BOLD responses of motion selective areas, and hence to facilitate their localization.

A forced binary decision task was used to estimate motion discrimination thresholds. Stimuli were presented in blocks of 64 trials. Each block lasted about 3 min. Each trial started with the presentation of the stimulus for 200 ms. Participants had to report whether the motion of the stimulus was upward or downward. They were instructed to respond as quickly as possible while maximizing their performances. Responses reported after 2 s were considered as incorrect. After each trial, an auditory feedback indicated whether the chosen direction was correct or not. During each block, we manipulated motion coherency (i.e., the percentage of dots moving along the same direction while the other dots had random directions) and estimated the thresholds corresponding to 80% of correct detection using an adaptive Bayesian approach (QUEST, see below). Participants completed 4 blocks. The first block was considered as a training and not included in the analyses. Breaks were included (3 minutes on average) between blocks to reduce fatigue. The whole experiment was completed in one session of about 20 minutes.

### 2.10 Robust estimation of motion coherence thresholds

Because it is complicated for MD patients to perform long psychophysical experiments, we used the QUEST adaptive procedure to obtain rapid, efficient and robust estimations of motion coherence thresholds (Guénot et al., 2022, 2023). QUEST is based on a Bayesian approach assuming that the psychophysics function underlying the participant performance follows a Weibull distribution. During a block, the estimated parameters of this function were updated after each trial on the basis of the participant’s response. Coherence values corresponded to the current maximum likelihood estimate of the threshold. We fixed the maximum number of trials at 64 as it was previously shown that this value leads to robust thresholds in most circumstances (Watson & Pelli, 1983). We used an initial threshold value of 58 % based on previous recordings using the same stimuli in MD patients (Guénot et al., 2022).

## 3. Results

In this section, we first detail the fMRI activation patterns observed in our MD patients and their controls. Next, we describe the results of the psychophysical experiment and their relationship with BOLD responses.

### 3.1 Motion-selective responses in the cortex of MD patients and their controls

Figure 3 shows the motion-selective activations (t-scores, p-value < 0.001, uncorrected) obtained in one typical MD patient (MD3, upper panel) and her control (CTRL3, lower panel) displayed on enlarged lateral views of their inflated individual cortical surfaces. Motion-selective activations in the other participants are shown in Figure 4. Activations displayed on medial views for all participants are shown on Figure 5. In all these figures, cortical nodes with significantly stronger responses to translational motion than to static dots appear in hot colors (yellow/red) while nodes with significantly stronger responses to the opposite contrast are shown in cold (blue) colors. Data were thresholded at p-value < 10^-3^ (uncorrected).

**Figure 5:**
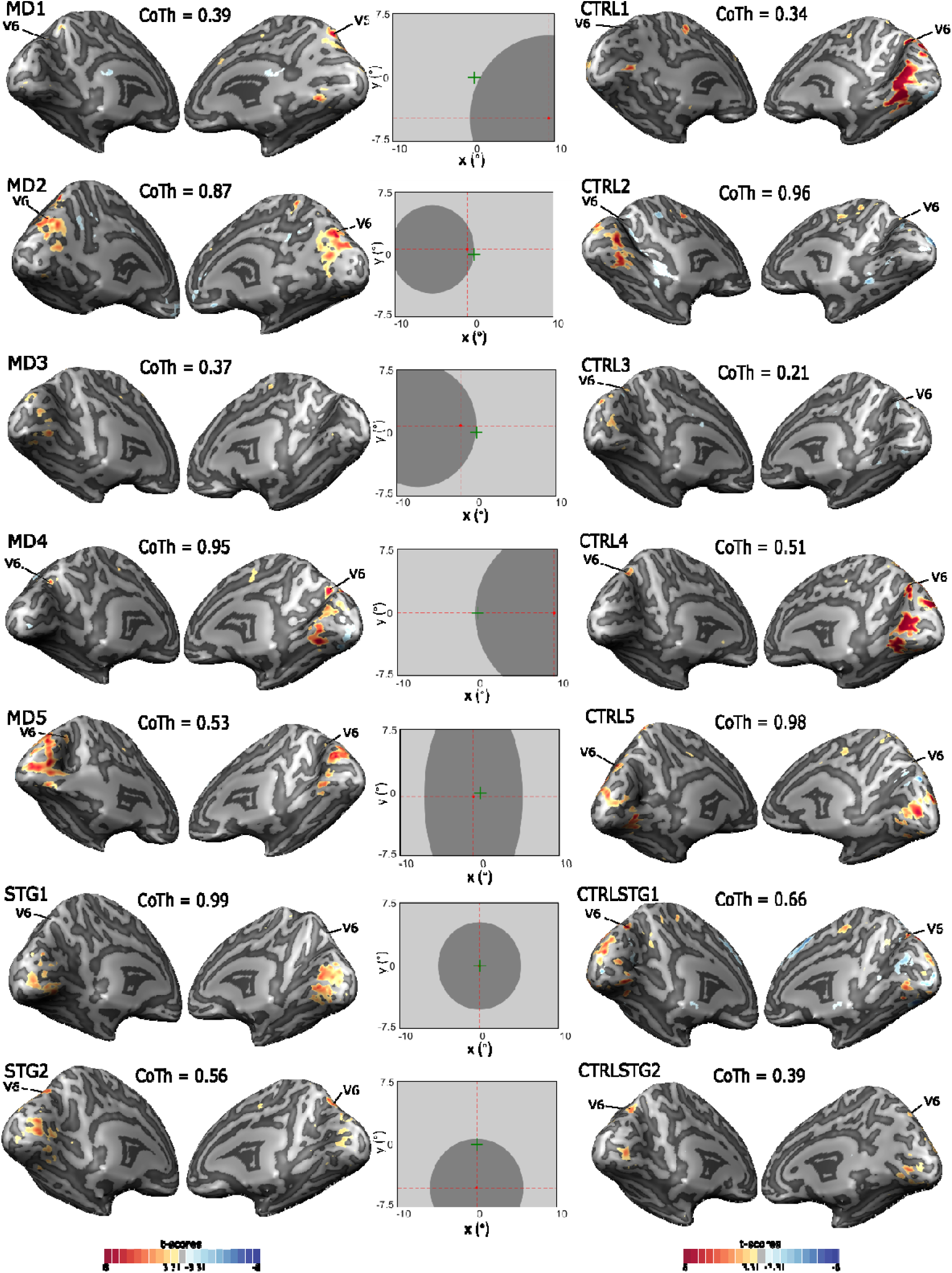
Comparison between BOLD responses to the translational motion condition and to its control condition (static dots) in all participants. Significantly stronger responses to motion are shown on medial views of the cortical surfaces for MD patients (leftward panels) and their controls (rightward panels). See Figure 3 for the details of the legend.

In all participants, translational motion elicited significantly stronger responses in a cortical network that is mainly localized within the dorsal visual pathway (i.e., in occipito-parietal areas) and within early visual areas. It should be noted that this network is very restricted when compared to the one obtained by contrasting responses to visual stimulation versus baseline (ocular fixation) and which covers the majority of the posterior part of the brain and notably includes areas along the ventral pathway (lateral and medial views of the corresponding activations maps are provided in Supplementary Figures 4-1 and 4-2). In the following, we first focus on responses within cortical regions that were consistently reported as motion-selective in the literature. Then, we describe our results in early visual areas and in other parts of the brain.

### 3.2 Activations in area hMT+

In all MD patients and their controls, we observed significantly stronger responses to translational motion than to static dots within (or posterior to) the ventral part of the anterior occipital sulcus (AOSv, in cyan in Figure 3), in between the inferior (IOS, in olive green) and the lateral (LOS, in green) occipital sulci (see Figures 3 and 4). The Talairach coordinates of the local maxima (t-scores) measured in this region are provided in Table 2. Averages values in the left and right hemispheres for patients ([-42±2; - 71±2; 5±2] and [41±2;-66±2; 2±2]) and controls ([-42±2;-64±2; 4±3] and [42±2;-63±2; 3±2]) were similar. These anatomical locations and coordinates are in very good agreement with those reported in the literature for area hMT+ (e.g., [-41;-69; 2] and [41;-67; 2] in Watson et al., 1993 or [-47;-76; 2] and [44;-67; 0] in Dumoulin, 2000). To confirm that our data reflected motion-selective activations in this area, we used independently defined regions of interest (ROIs) centered on the Talairach coordinates provided by Watson et al., (1993) for hMT+ and tested whether the differences between the percentages of signal change (PSCs) measured during the translational motion condition and during its control (static dots) were significantly different from zeros across participants in this ROI (see the ‘*Methods*’ section). This was the case for both groups (Wilcoxon sign-rank tests, p-value = 0.0156 in MD patients and p-value = 0.0078 in controls, see Figure 6-A). Importantly, we did not find significant differences between the differences in PSCs observed in this ROI for the two groups (Wilcoxon paired sign-rank tests, p-value = 0.1094, Bayes factor = 1.6). As an additional analysis, we also considered the activation extent within this ROI and found that they did not significantly differ between the two groups (see Supplementary figure 6-1). Altogether, these results suggest that both MD patients and their controls have motion-selective responses in area hMT+ and that these responses do not significantly differ between the two populations.

**Figure 6:**
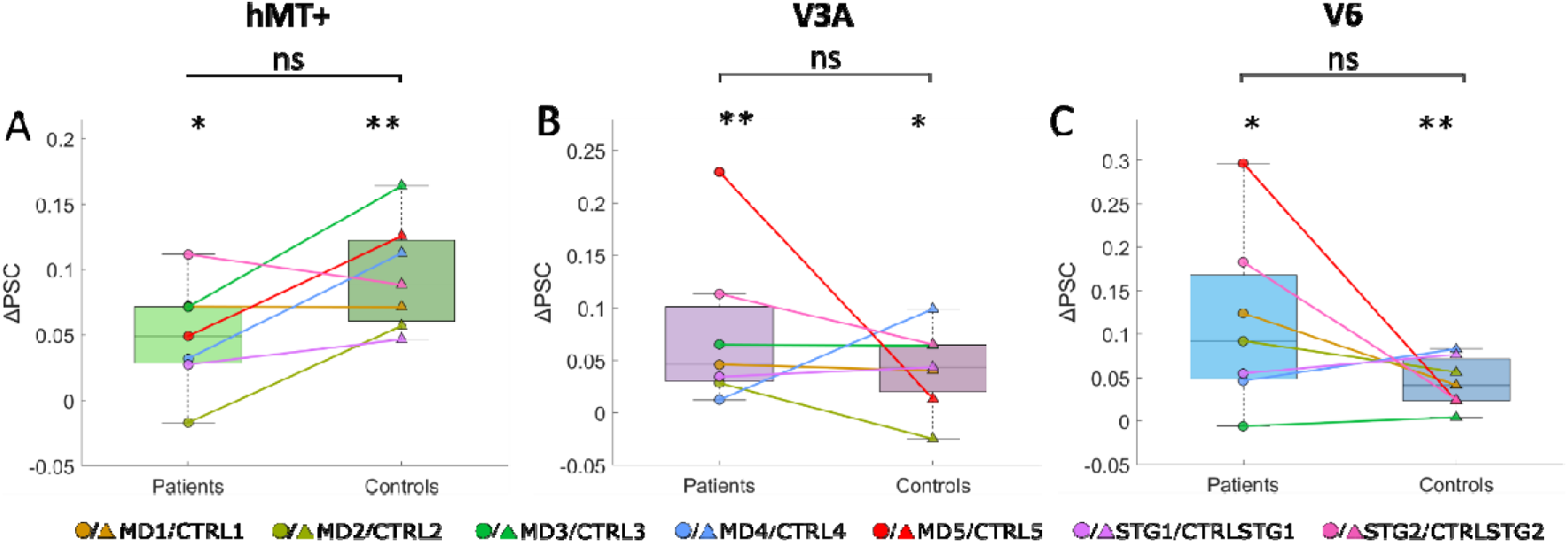
Motion-selective activations in the hMT+ (A), V3A (B) and V6 (C) ROIs. For each patient and their age-matched control (connected by lines), colored dots provide the differences between the average (across voxels and hemispheres) percentages of signal changes (PSCs) during the translational motion and control (static dots) conditions. For each group, the boxplots provide the first quartile, median, last quartile, and the extreme PSC difference. Stars (* or **) indicate that the PSC difference is significantly different from zero (p-values < 0.05 or <0.01, respectively). Statistical analyses based on paired Wilcoxon sign-rank tests did not reveal significant differences between activations in the two groups (p-values > 0.05 for the three ROIs).

**Table 2:**
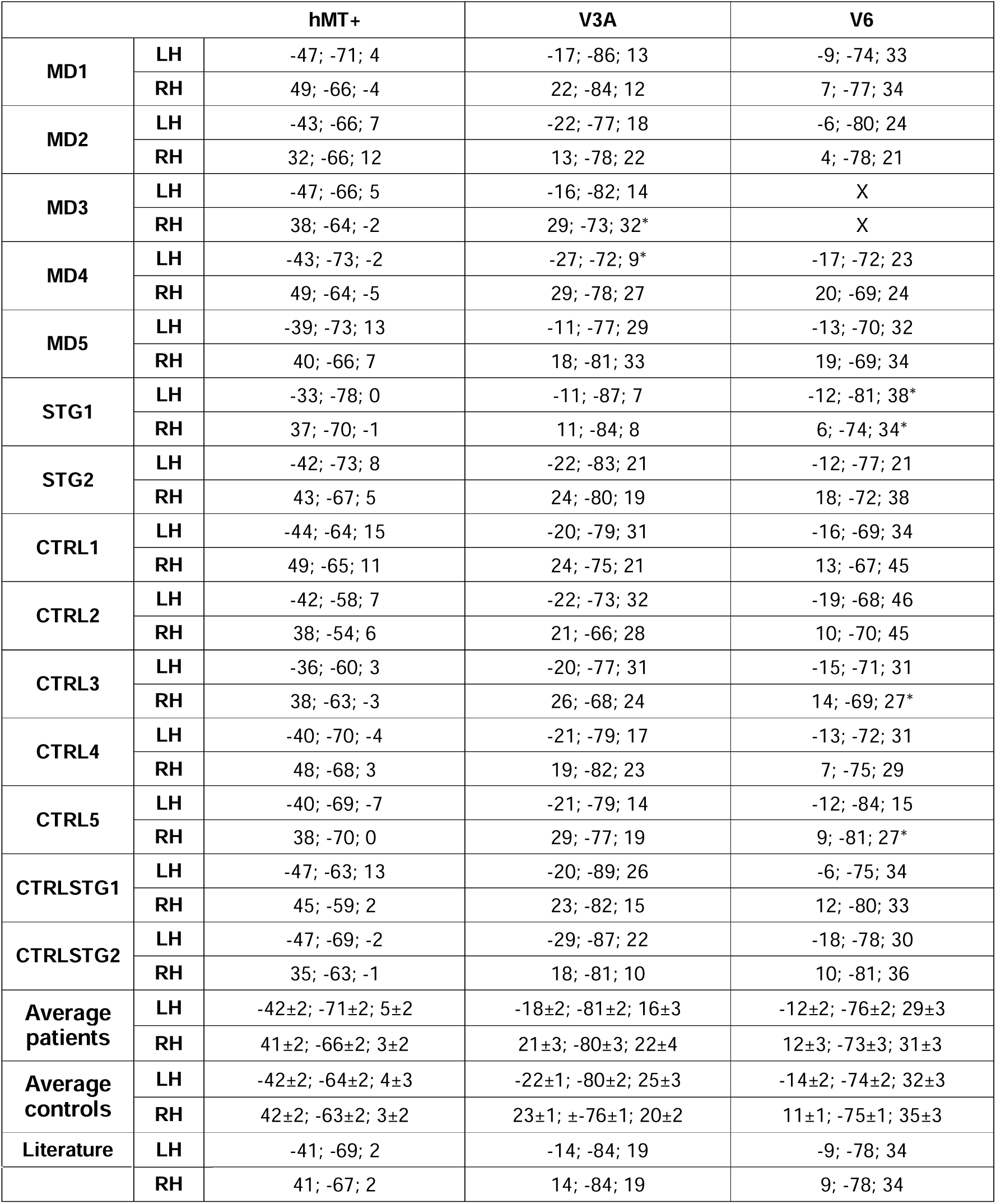
Talairach coordinates of the different motion-selective areas observed in our study (hMT+, V3A and V6). Coordinates are provided for each MD patient and his/her control. The ‘*’ symbol indicates the coordinates for which a more permissive threshold has been used, i.e p<10-2. The table also provides average coordinates across each group as well as average coordinates reported in previous studies for these 3 areas (Watson et al., 1993 for hMT+; Tootell et al., 1997 for V3A and Pitzalis et al., 2010 for V6).

|  |  | hMT+ | V3A | V6 |
| --- | --- | --- | --- | --- |
| MD1 | LH | -47; -71; 4 | -17; -86; 13 | -9; -74; 33 |
|  | RH | 49; -66; -4 | 22; -84; 12 | 7; -77; 34 |
| MD2 | LH | -43; -66; 7 | -22; -77; 18 | -6; -80; 24 |
|  | RH | 32; -66; 12 | 13; -78; 22 | 4; -78; 21 |
| MD3 | LH | -47; -66; 5 | -16; -82; 14 | X |
|  | RH | 38; -64; -2 | 29; -73; 32* | X |
| MD4 | LH | -43; -73; -2 | -27; -72; 9* | -17; -72; 23 |
|  | RH | 49; -64; -5 | 29; -78; 27 | 20; -69; 24 |
| MD5 | LH | -39; -73; 13 | -11; -77; 29 | -13; -70; 32 |
|  | RH | 40; -66; 7 | 18; -81; 33 | 19; -69; 34 |
| STG1 | LH | -33; -78; 0 | -11; -87; 7 | -12; -81; 38* |
|  | RH | 37; -70; -1 | 11; -84; 8 | 6; -74; 34* |
| STG2 | LH | -42; -73; 8 | -22; -83; 21 | -12; -77; 21 |
|  | RH | 43; -67; 5 | 24; -80; 19 | 18; -72; 38 |
| CTRL1 | LH | -44; -64; 15 | -20; -79; 31 | -16; -69; 34 |
|  | RH | 49; -65; 11 | 24; -75; 21 | 13; -67; 45 |
| CTRL2 | LH | -42; -58; 7 | -22; -73; 32 | -19; -68; 46 |
|  | RH | 38; -54; 6 | 21; -66; 28 | 10; -70; 45 |
| CTRL3 | LH | -36; -60; 3 | -20; -77; 31 | -15; -71; 31 |
|  | RH | 38; -63; -3 | 26; -68; 24 | 14; -69; 27* |
| CTRL4 | LH | -40; -70; -4 | -21; -79; 17 | -13; -72; 31 |
|  | RH | 48; -68; 3 | 19; -82; 23 | 7; -75; 29 |
| CTRL5 | LH | -40; -69; -7 | -21; -79; 14 | -12; -84; 15 |
|  | RH | 38; -70; 0 | 29; -77; 19 | 9; -81; 27* |
| CTRLSTG1 | LH | -47; -63; 13 | -20; -89; 26 | -6; -75; 34 |
|  | RH | 45; -59; 2 | 23; -82; 15 | 12; -80; 33 |
| CTRLSTG2 | LH | -47; -69; -2 | -29; -87; 22 | -18; -78; 30 |
|  | RH | 35; -63; -1 | 18; -81; 10 | 10; -81; 36 |
| Average patients | LH | -42±2; -71±2; 5±2 | -18±2; -81±2; 16±3 | -12±2; -76±2; 29±3 |
|  | RH | 41±2; -66±2; 3±2 | 21±3; -80±3; 22±4 | 12±3; -73±3; 31±3 |
| Average controls | LH | -42±2; -64±2; 4±3 | -22±1; -80±2; 25±3 | -14±2; -74±2; 32±3 |
|  | RH | 42±2; -63±2; 3±2 | 23±1; ±-76±1; 20±2 | 11±1; -75±1; 35±3 |
| Literature | LH | -41; -69; 2 | -14; -84; 19 | -9; -78; 34 |

|  | RH | 41; -67; 2 | 14; -84; 19 | 9; -78; 34 |
| --- | --- | --- | --- | --- |

### 3.3 Activations in area V3A

In all MD patients and their controls, we observed significantly stronger responses to translational motion within the transverse occipital sulcus (in purple on Figure 3, see also Figure 4). For some participants, we had to decrease our threshold to a p-value of 10^-2^, uncorrected) at similar Talairach coordinates between the two groups ([-18±2-81±2 16±3] and [21±3-80±2 22±4] for patients and [-22±2-80±1 25±3] and [23±1-76±1 20±2] for controls on average (see Table 2). These anatomical locations and coordinates are in very good agreement with those reported in the literature for area V3A (e.g., [±14;-84; 19] in Tootell et al., 1997 or [±18;-78; 25] in Pitzalis et al., 2006). For both groups, we confirmed that our data reflected motion-selective activations in this area from statistical analysis within an ROI centered on the Talairach coordinates provided by Tootell et al. (1997) (Wilcoxon sign-rank tests, p-value = 0.0078 in MD patients and p-value = 0.0234 in controls, see Figure 6-B and above). We did not find significant differences between the differences of PSCs observed in this ROI for the two groups (Wilcoxon paired sign-rank tests, p-value = 0.469, Bayes factor = 0.5), nor between their activation extents (see Supplementary figure 6-1). Altogether, these results suggest that both MD patients and their controls have motion-selective responses in area V3A and that these responses do not significantly differ between the two populations.

### 3.4 Activations in area V6

In six MD patients and their controls (all participants except MD3 and CTRL3), we observed significantly stronger responses to translational motion in the upper part of the cuneus, within the posterior branch of the parieto-occipital sulcus (POS) at similar Talairach coordinates between the two groups ([-12±2-76±2 29±3] and [12±3 - 73±3 31±3] for patients and [-14±2-74±2 32±3] and [11±1-75±1 35±3] for controls on average, see Table 2). These anatomical locations and coordinates are in very good agreement with those reported in the literature for area V6 (e.g., [±9;-78; 34] in Pitzalis et al., 2010). For both groups, we confirmed that our data reflected selectivity to translational motion in this area from statistical analysis within an ROI centered on the Talairach coordinates provided by Pitzalis et al. (2010) (Wilcoxon sign-rank tests, p-value = 0.0156 in MD patients and p-value = 0.00781 in controls, see Figure 6-B and above). We did not find significant differences between the differences of PSCs observed in this ROI for the two groups (Wilcoxon paired sign-rank tests, p-value = 0.297, Bayes factor = 0.7), nor between their activation extents (see Supplementary figure 6-1). Altogether, these results suggest that both MD patients and their controls have motion-selective responses in area V6 and that these responses do not significantly differ between the two populations.

### 3.5 Activations in early visual areas and relationship to retinotopic properties

In all participants, we also observed motions selective activations below and above the most anterior part of the calcarine sulcus (see Figure 5). These regions correspond to the portions of primary visual areas (V1, V2 and V3) connected to peripheral vision. Their stronger responses to translational motion are not surprising given that a vast proportion of their neurons are motion-direction selective (Dumoulin et al., 2003). Here as well, we did not observe different patterns of activations in statistical parametric maps between groups (Figure 5). It should be noted that for patients whose scotoma covers a larger surface in one visual hemifield (because their PRL is lateralized) and for their controls (MD1/CTRL1, MD2/CTRL2, MD3/CTRL3 and MD4/CTRL4), motion-selective activations were generally stronger in the hemisphere ipsilateral to the scotoma (i.e., contralateral to the visual hemifield that receives more visual inputs) which reflects the retinotopic properties of early visual areas (see e.g., Benson & Winawer, 2018). In V3A, hemispheric asymmetries were also observed for these participants but they were less pronounced, reflecting the fact that neural receptive fields in this area are bigger than in areas V1, V2 and V3 (e.g., Barton & Brewer, 2017 reported population receptive fields with radiuses between 4 and 6° at eccentricities comparable to those in our study). We did not observe hemispheric asymmetries in areas hMT+ and V6, probably because these areas have very large receptive fields which overlap with the ipsilateral visual hemifield (for hMT+, see Huk et al., 2002; for V6, see Pitzalis et al., 2010). For hMT+, this is notably true in its hMST functional subdivision.

### 3.6 Additional activations

Beyond the activations reported above, we also observed motion-selective responses in other parts of the brain but they were not as reliable and robust across participants. In some patients and their controls (MD2/CTRL2, MD3/CTRL3, MD4/CTRL4, STG1/CTRLSTG1 and STG2/CTRLSTG2, see also on the right hemisphere of CTRL5), we notably observed motion-selective activations in the fundus of the superior temporal sulcus (STS, in red on Figure 3) and also (but less frequently) on its lower bank (MD3 in the right hemisphere, MD4 in the right hemisphere, CTRL5 in the left hemisphere) or within the middle temporal sulcus (MTS, CTRL2). These activations could correspond to the human posterior temporal sulcus (pSTS) area, which is selective to biological motion (Grossman & Blake, 2002) and which preferentially encodes peripheral vision (Barton & Brewer, 2017). The associated Talairach coordinates are in good agreement with those reported for this area in the literature (see supplementary Table 2-1) but an ROI-based analysis using a sphere centered on the coordinates provided by Barton and Brewer (2017) did not confirm its implication (Wilcoxon sign-rank tests, p-value = 0.109 in both MD patients and controls, see Supplementary Figure 6-2).

In some participants (MD3, MD4, CTRL1 and CTRL2), additional motion-selective activations were also observed in the parietal cortex and notably within the intra-parietal sulcus, where is located the ventral intraparietal (VIP) area, which is motion selective (Bremmer et al., 2001).

We saw that two patients (MD5 and STG1) relied on central vision for fixation, indicating that they retained some residual visual function near their fovea (see the ‘Methods’). To ensure that their inclusion did not bias our results, we replicated all statistical analyses described above excluding these patients (and their matched controls) and found consistent effects (see Supplementary Figure 6-3).

### 3.7 Motion-direction discrimination thresholds and their relationship with BOLD responses

The motion direction discrimination thresholds estimated in each MD patient and their age-matched control (see the ‘Material and Methods’ section) are provided in Figure 7-A. Thresholds varied between 0.37 and 0.99 in patients (0.37 for MD3 whose fMRI data are also shown on Figure 3; 0.67 on average) and between 0.21 and 0.98 in control participants (0.21 for CTRL3, see Figure 3; 0.58 on average). We did not find significant differences between the thresholds measured in the two groups (Wilcoxon signed-rank test, p-value = 0.1698, Bayes factor = 0.25). Altogether, these results confirm Guénot et al.’s (2022) results where motion direction discrimination thresholds did not significantly differ between MD patients and their age-matched controls under identical experimental conditions (see Figure 5 in Guénot et al., 2022).

**Figure 7:**
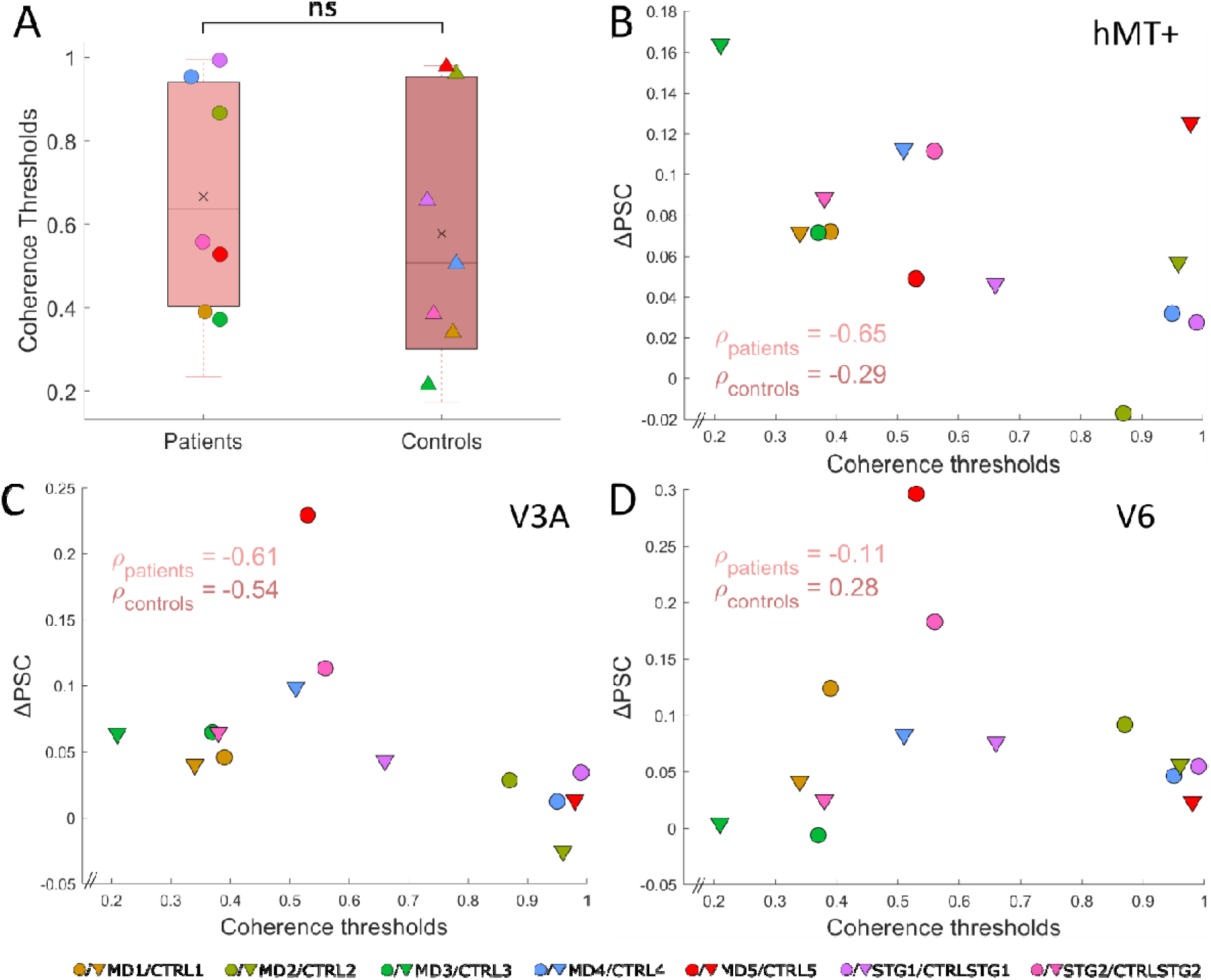
Behavioral results and relationship with cortical responses in motion-selective ROIs. A) Motion-direction discrimination thresholds estimated in the two groups. The boxplots show the different values for the first quartile, median, last quartile, and the extreme data points. The cross represents the means for both groups. Dots provide the individual data points of the distributions. It should be noted that they were slightly offset horizontally to improve their visibility. B) Differences between the average percentages of signal changes (PSCs) during the translational motion and control (static dots) conditions in the hMT+ ROI as a function of coherence thresholds. Dots and triangles respectively correspond to MD patients and controls. Spearman correlation coefficients rho (⍴) are shown (the associated p-values were respectively 0.1389 for patients and 0.5560 for controls). Data in the V3A and V6 ROIs are respectively provided in panels C and D. The p-values associated with the Spearman correlation coefficients were 0.1667 and 0.2357 in area V3A and 0.8397 and 0.5560 in area V6 for patients and control participants, respectively. The x-axis was cropped at 0.2 for better visibility.

Next, we examine whether the performances measured in psychophysics are correlated with fMRI activations in motion-selective areas. For each participant, panels B, C and D of Figure 7 show the differences between the averages percentages of signal changes (PSCs) during the translational motion condition and during its control condition (static dots) in areas hMT+, V3A and V6 as a function of motion-discrimination thresholds. For each group, we provide the corresponding Spearman correlation coefficients. This measure tests whether the data are monotonically related and do not assume normality. Although stronger responses to motion stimuli were generally associated with lower perceptual thresholds (Spearman coefficients were mostly negative), we did not observe any significant correlations in the data (p-values > 0.5, see details in the legend of figure 7).

To complete these analyses, we also tested whether the size of the scotoma could influence the percentages of signal changes measured in our ROIs but we did not observe significant effects in both groups (see Supplementary Figure 7-1). This is probably because receptive fields in these areas are large enough to capture visual inputs outside the scotoma (see also the **‘***Activations in early visual areas and relationship to retinotopic properties*’ section).

### 3.8 Control analyses to verify the robustness of our ROI-based approach

All the ROIs-based analyses in our study used spheres of 10 mm. To make sure that this choice did not impact our results, we reproduced these analyses with spheres of 5 and 15 mm of radius and found the same patterns of results.

## 4. Discussion

The aim of this study was to characterize the cortical network processing translational motion in maculopathy patients. We recorded fMRI data in 7 patients and in 7 age-and gender-matched controls using an experimental paradigm based on RDKs classically employed to identify motion selective areas (Zeki et al., 1991; Watson et al., 1993; Tootell et al., 1995a; Huk et al., 2002). We also estimated motion-direction discrimination thresholds in each participant. Stimuli in each control participant were masked using an artificial scotoma of the same dimension as the scotoma of their paired patients, in order to equalize the visual inputs received by the two populations.

A previous study on an animal model (Burnat et al., 2017) demonstrated that inducing a central retinal lesion in adult cats enhanced neuronal activity in the posteromedial lateral suprasylvian area (PMLS), the homologue of primate cortical area V5/MT (Villeneuse et al., 2006), and to a lesser extent in area 7, which is also motion-sensitive (Pigarev & Rodionova, 1998). Using a population receptive field analysis, Shao et al. (2013) found that fMRI activations in area V5/MT were more extensive in a macaque with juvenile MD compared to control monkeys with a 10° artificial scotoma. Additionally, a recent human study revealed enhanced functional connectivity between MT and early visual areas responsive to peripheral vision in individuals with central vision loss (Fleming et al., 2024). Based on these findings, we hypothesized that the cortical network processing motion might reorganize in MD patients in the presence of the scotoma through enhanced BOLD signals in motion-selective areas and eventually through an expansion of these areas. However, our observations indicated that the cortical networks activated by motion were similar between MD patients and controls, although with only 2 patients suffering from late-onset Stargardt disease, we cannot make a definitive statement here. In both populations, our protocol elicited significant and robust responses in areas hMT+, V3A and V6, as well as within the portion of early visual areas connected to peripheral vision. An ROI-based analysis confirmed the implication of these regions and the associated Talairach coordinates are in good agreement with those reported in the literature. Importantly, we did not observe significant motion-selective activations outside of this network, which thus rules out the possibility that new motion areas emerged in patients after the disease apparition. Furthermore, we found no significant differences in BOLD response amplitudes and extents between patients and controls in hMT+, V3A, and V6. In V3A and V6, the null hypothesis was supported by Bayes factors (<1), although a larger sample is needed for firmer conclusions. In hMT+, a slight trend toward weaker and less extensive activations in patients compared to controls was noted (Bayes factors = 1.6 and 1.5), but this trend was not significant and actually goes against our hypotheses about reorganization. Overall, our results do not indicate any significant large-scale enhancement of the motion-selective network in MD patients. It is possible that the effects observed by Burnat et al. (2017) in cats (and also in macaques by Shao et al., 2013) are not transferable to humans. Indeed, it has been suggested that retinal damage in MD may trigger cortical degeneration in high-level areas (Hernowo et al., 2014). This raises the possibility that some of the motion-selective regions studied here could be affected by atrophic processes secondary to sensory deprivation, limiting their capacity to reorganize. Alternatively, maybe reorganizations exist in humans but task-based fMRI is not an appropriate neuroimaging technique to capture them since it only provides an indirect measure of cortical activity (see Calford et al., 2005).

From their fMRI recordings performed in macaques, Shao et al. (2013) also found that population receptive field (pRF) size in area V5/MT was significantly smaller in an animal with juvenile MD. To better understand whether this effect is also present in humans, it will be important to reproduce retinotopic mapping experiments in MD patients and to perform a pRF analysis. Although several research groups demonstrated that such an approach is feasible in patients suffering from different kinds of visual pathology (Carvalho et al., 2022; Papanikolaou et al., 2014; Silson et al., 2023), it remains a challenge in maculopathy patients because their eye fixation is usually not stable enough to obtain reliable signals.

While numerous neuroimaging studies on cortical reorganization in MD patients focused on primary visual areas (see e.g., Morland, 2015 for a complete review), much less investigated how activations in higher-level cortical areas are modified in maculopathy. Little et al. (2008) found that age-related MD patients had stronger BOLD responses than control participants in the prefrontal cortex and in the intraparietal sulcus during visually guided saccade and smooth-pursuit tasks, two cognitive functions that rely on a form of motion processing. Important modifications in brain activity were also observed in MD patients during word recognition (Szlyk & Little, 2009) and scene processing (Ramanoël et al., 2018), maybe because the visual field in control participants was not masked by an artificial scotoma. It was suggested that enhanced fMRI activations measured in higher-level cortical areas in these cases reflected a form of compensation for the decreased sensory functions. If we did not observe significant modifications of brain activity in our case, it might be because motion perception in MD patients is not significantly impaired in our task and no compensation is thus needed.

We estimated motion direction discrimination thresholds in all our participants because we wanted to replicate one previous result from Guénot et al. (2022) and also to determine whether fMRI activations in some cortical areas were correlated with motion perception. We found that discrimination thresholds for lateral motion do not significantly differ between patients and controls under similar visual conditions. This confirms and further supports the results of Guénot et al. (2022) which were obtained using the same protocol but with other groups of patients and controls. While TMS studies suggested that hMT+ is causally involved in motion perception (Chakraborty et al., 2021; McKeefry et al., 2008; Strong et al., 2017) and fMRI activations in this area under different motion conditions were shown to correlate with behavioral reports (Tootell et al., 1995), we did not observe significant correlations between BOLD activations in our different motion-selective areas (notably hMT+, V3A and V6) and perceptual thresholds. It should be noted that in our case, correlations were estimated across MD patients (or across controls), while Tootell et al. (1995b) examined intra-individual correlations across experimental conditions. The important inter-individual variability observed in motion-discrimination thresholds and in brain activations in our data as well as our limited number of participants might explain why our results differ. Another explanation could also lie in the difference between the protocols used during our psychophysical and fMRI experiments. Indeed, in order to maximize BOLD responses, our stimuli alternated between dots moving coherently during 1.8s (with an unlimited lifetime) and static dots under the MRI scanner while our behavioural protocol involved dots moving for 0.2s with a limited lifetime. It was reported that small changes in paradigm based on random dots can influence the results (Pilly & Seitz, 2009).

A limitation of our study is the limited number of participants (7 patients and their controls). Given this small sample size and our methodology specifically designed to be suitable in this case, it should be viewed as a case study with matched controls. It is very challenging to recruit MD patients for neuroimaging experiments and this number can be explained by the inclusion criteria that we applied. We only recruited patients with no long-term illnesses, with a stable PRL on at least one eye, with typical cognitive functions and psychological states (as evaluated by the MMSE and BDI-II), and with no contraindication for fMRI recordings. The present study is nonetheless in line with previous neuroimaging works in maculopathy patients (e.g., 4 patients were scanned in Ramanoël et al., 2018, 6 in Little et al., 2008, 8 in Szlyk & Little, 2009 and 6 in Masuda et al., 2021). Because of our small number of participants, our analyses lack the statistical power to definitively rule out all forms of reorganization in MD patients (notably, our Bayes factors only indicate a weak evidence against enhanced responses in patients). The reliable activation maps that we obtained in all our patients and their controls (with no significant differences between the response strengths and extent observed within motion selective areas in the two populations) nonetheless suggest that the motion processing network is preserved and does not undergo large-scale enhancements in MD patients following the onset of the scotoma.

We did not perform whole brain analyses because there was an important variability between the scotomas and PRL positions of our MD patients and hence between their visual inputs and between the associated cortical activations. It makes it difficult to obtain reliable results once fMRI responses across patients (or across controls) are projected onto a global brain template. This problem is further amplified by the important variability that exists between the position and extent of functional visual areas across individuals (see e.g., Kolster et al., 2010 for the hMT+ cluster). We rather chose to analyze the data at the individual level and to characterize the motion-selective network in each patient and their age-matched controls. We performed group-level pairwise statistics using independently defined regions-of-interest (ROIs), which permits us to get rid of the multiple comparisons problem (see e.g., Poldrack, 2007).

## Acknowledgments

This study was supported by a French grant from the Agence Nationale de la Recherche (ANR-21-CE28-0021, ANR PRC ReViS-MD; awarded to C.P. and B.R.C.). This work was performed on the Inserm/UPS UMR1214 Technical Platform of Toulouse and the IRMaGe platform member of France Life Imaging network (grant ANR-11-INBS-0006) for Grenoble. We thank Judith Eck for helping on fMRI analysis, and Ilia Korjoukov and Louise Kauffmann for helping develop our MRI paradigm.

## Supplementary Materials

**Supplementary Figure 1-1:**
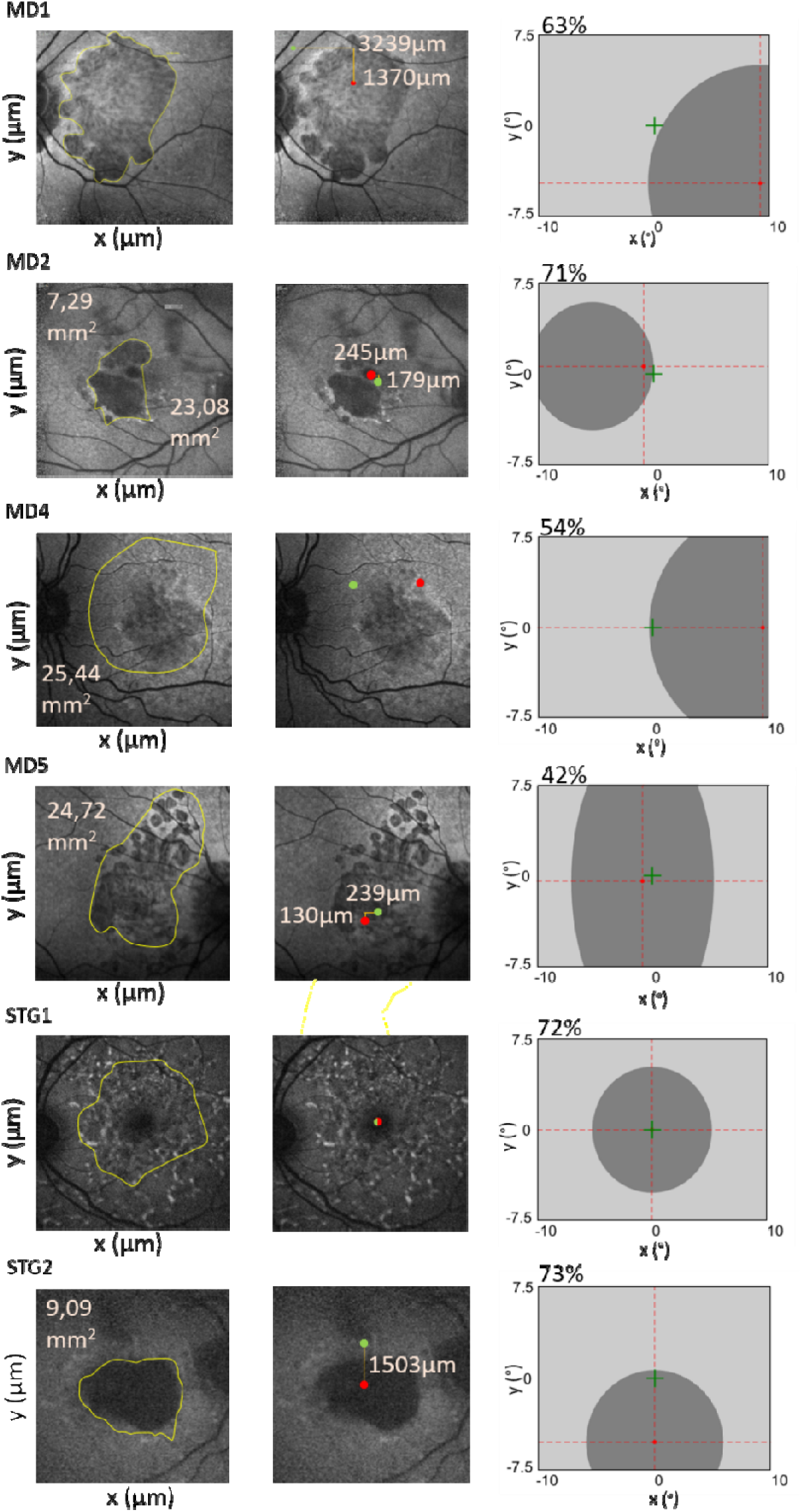
Images of the eye fundus and estimation of the scotoma size (left panel) and PRL (middle panel) in all patients (except MD3, whose data are provided in figures 1 and 3). The right panels show the artificial scotomas used in the corresponding control participants. In each case, we provide the visible portion of the stimulus (i.e., the surface that is not covered by the artificial scotoma). It is to be noted that the fovea and the PRL are located at the same place for STG1. See Figure 1 for more details.

**Supplementary Figure 4-1:**
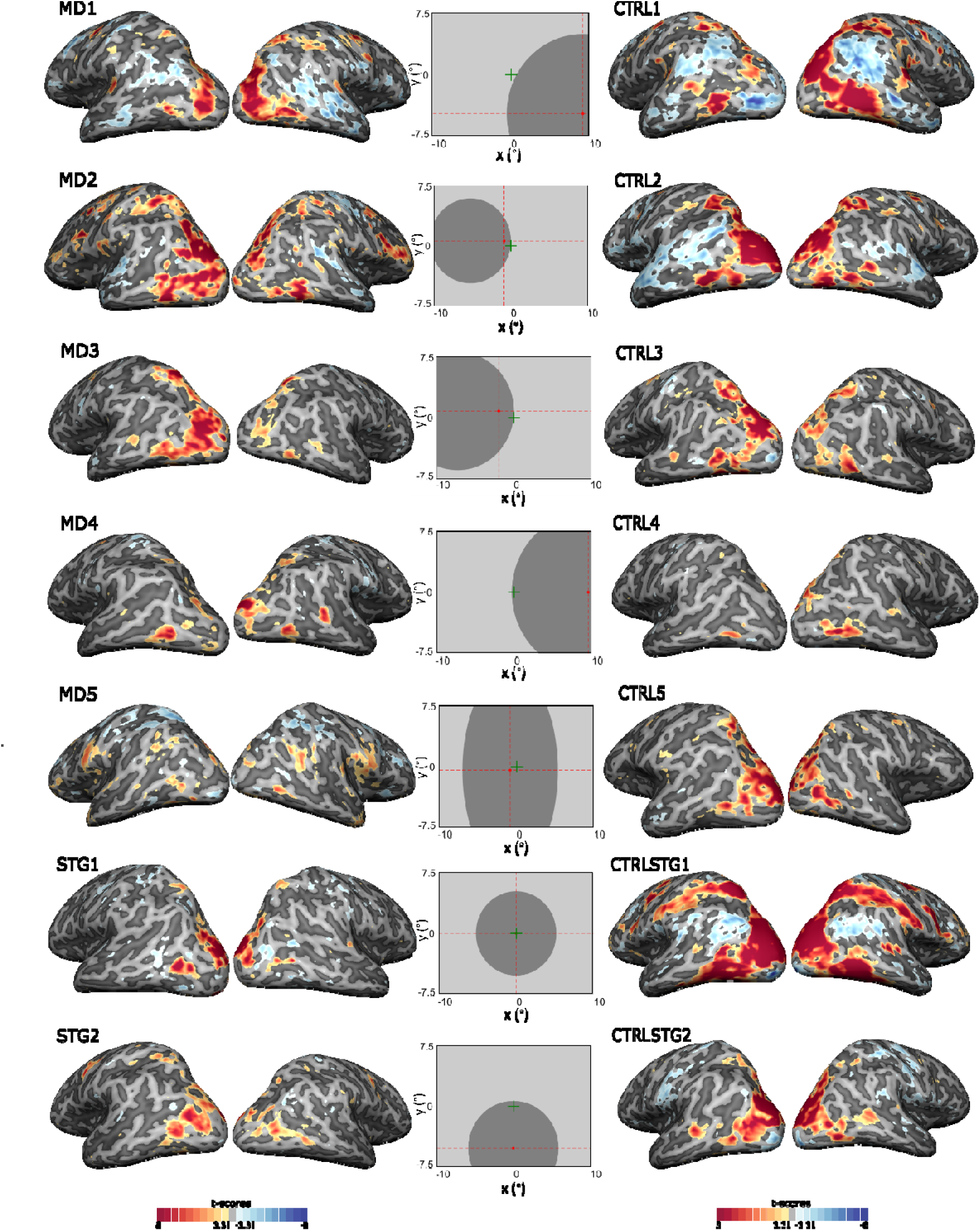
Comparison between BOLD responses to visual stimulation and to a blank screen on the lateral cortical surfaces of all MD patients (left panels) and their respective controls (right panels). See Figure 3 for the details of the legend.

**Supplementary Figure 4-2:**
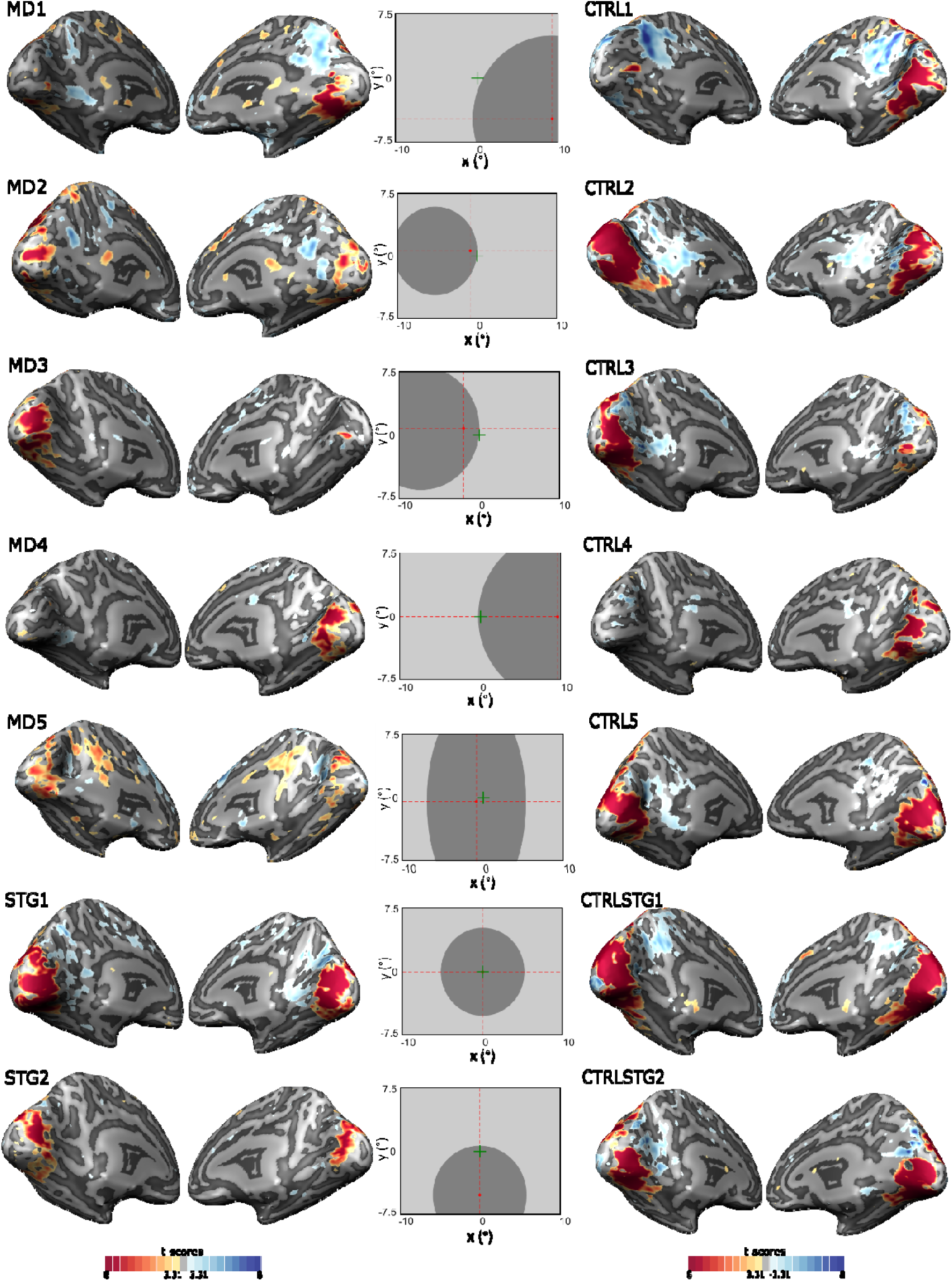
Comparison between BOLD responses to visual stimulation and to a blank screen on the medial cortical surfaces of all MD patients (left panels) and their respective controls (right panels). See Figure 3 for the details of the legend.

**Supplementary Figure 6-1:**
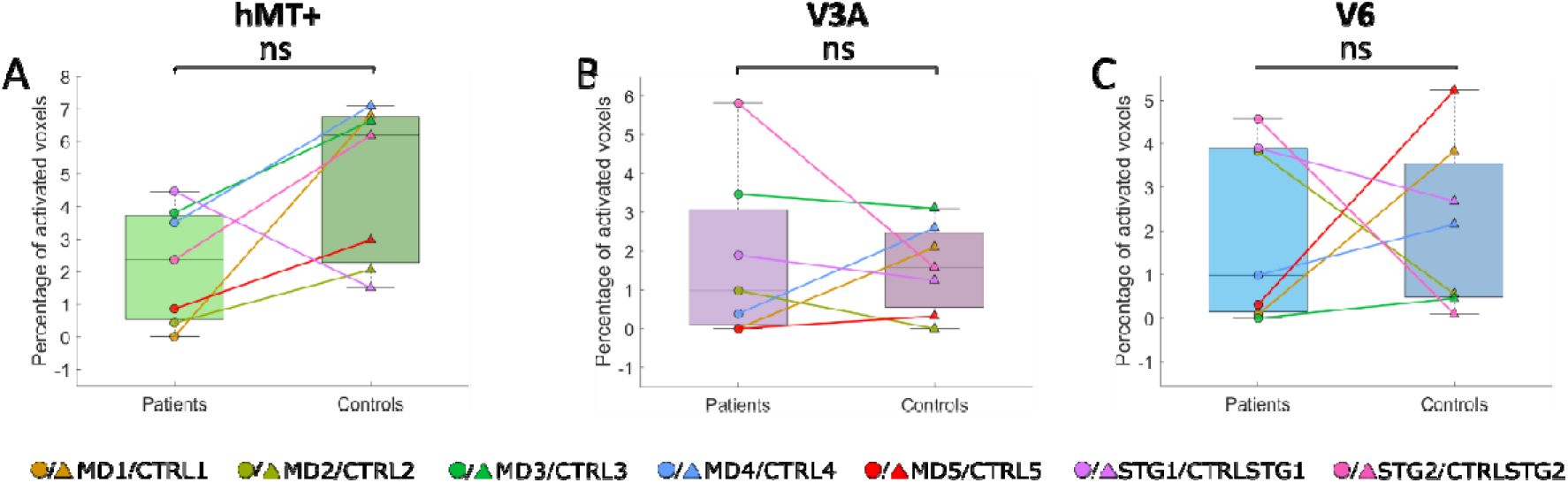
Activation extents with the hMT+ (A), V3A (B) and V6 (C) ROIs. These extents were computed as the percentage of voxels passing a threshold of t-score = 3 within each ROI. For each group, the boxplots provide the first quartile, median, last quartile, and the extreme percentages of activated voxels. None of the comparison led to significant results for either hMT+ (p-value > 0.05, Bayes factor = 1.5), V3A (p-value > 0.05, Bayes factor = 0.4) or V6 (p-value > 0.05, Bayes factor = 0.4). ROIs were defined as spheres of 10 mm of radius centered on the Talairach coordinates provided in the literature (see the ‘*Methods*’ section) but similar results were obtained when considering spheres of 5 and 15 mm of radius.

**Supplementary Figure 6-2:**
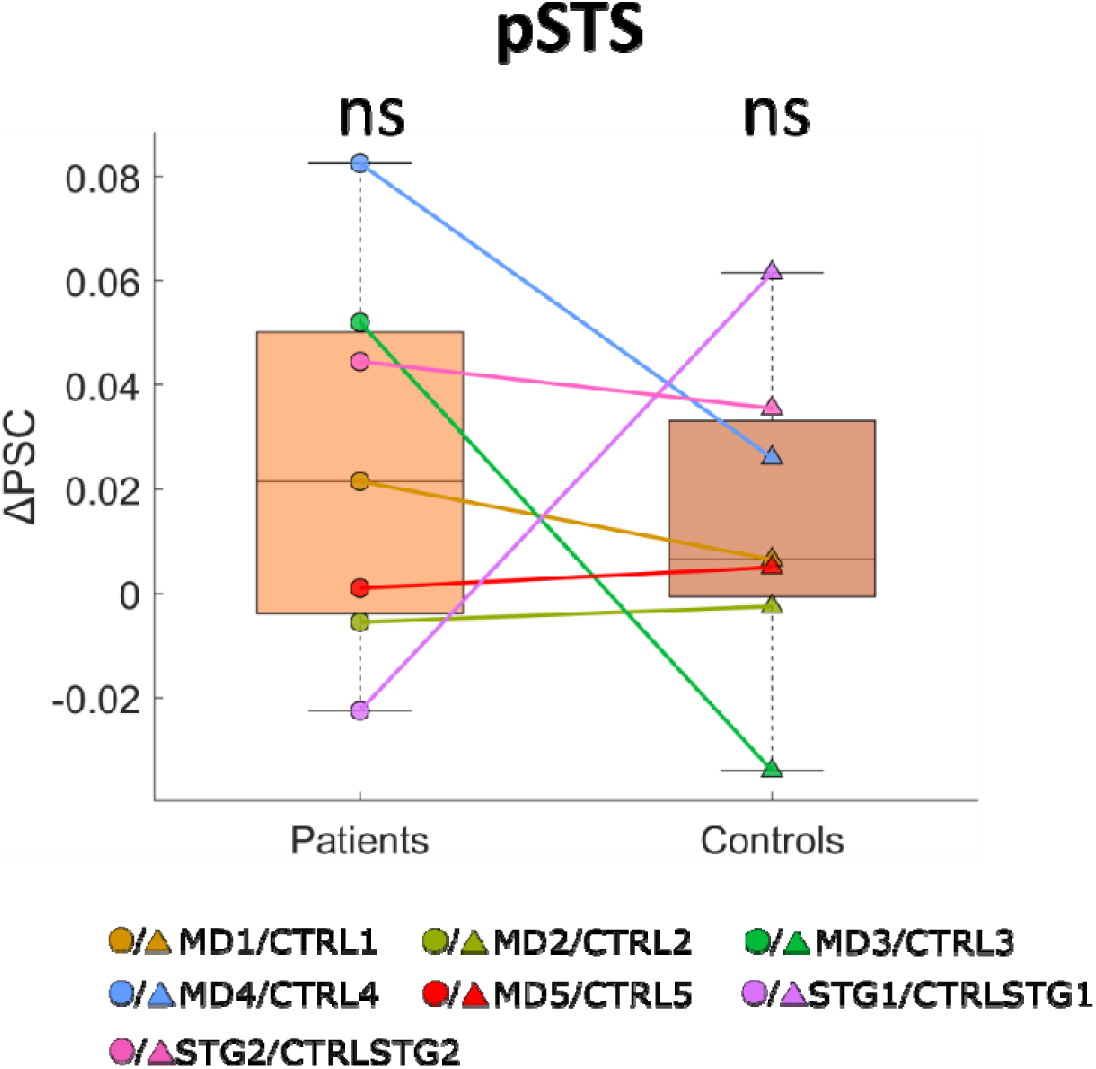
Motion-selective activations in the pSTS ROI. For each patient and their age-matched control (connected by lines), colored dots provide the differences between the average (across voxels and hemispheres) percentages of signal changes (PSCs) during the translational motion and control (static dots) conditions. For each group, the boxplots provide the first quartile, median, last quartile, and the extreme PSC difference. These differences are not significantly different from zero (p-values = 0.109 in both MD patients and controls, Wilcoxon test). Statistical analyses based on paired Wilcoxon sign-rank tests did not reveal significant differences between activations in the two groups (p-value > 0.05, Bayes factor = 0.47).

**Supplementary Figure 6-3:**
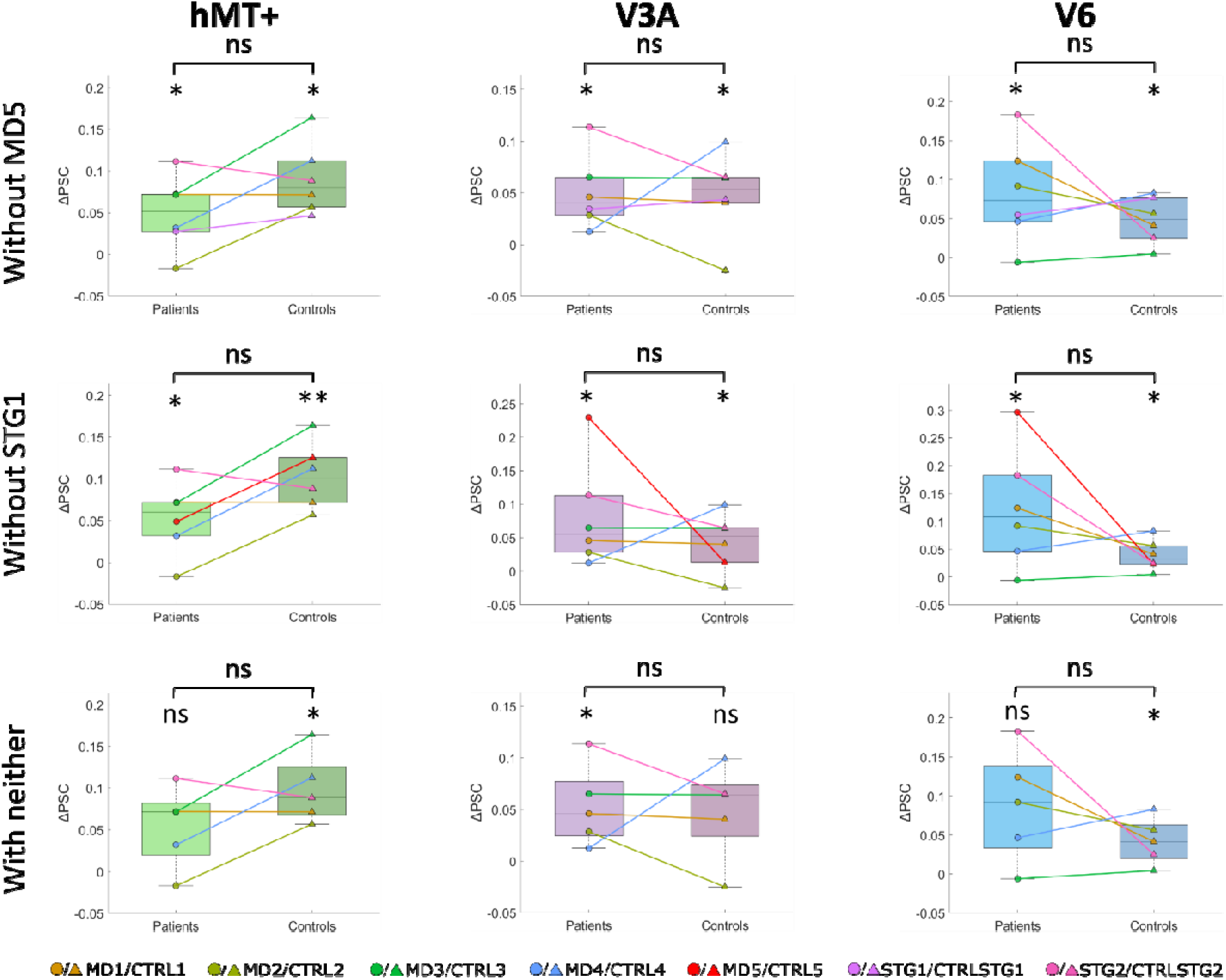
Motion-selective activations in the hMT+ (A), V3A (B) and V6 (C) ROIs when removing MD5, STG1, or both. For each patient and their age-matched control (connected by lines), colored dots provide the differences between the average (across voxels and hemispheres) percentages of signal changes (PSCs) during the translational motion and control (static dots) conditions. For each group, the boxplots provide the first quartile, median, last quartile, and the extreme PSC difference. Stars (* or **) indicate that the PSC difference is significantly different from zero (p-values < 0.05 or <0.01, respectively). Statistical analyses based on paired Wilcoxon sign-rank tests did not reveal significant differences between activations in the two groups (p-values > 0.05 for the three ROIs).

**Supplementary Figure 7-1:**
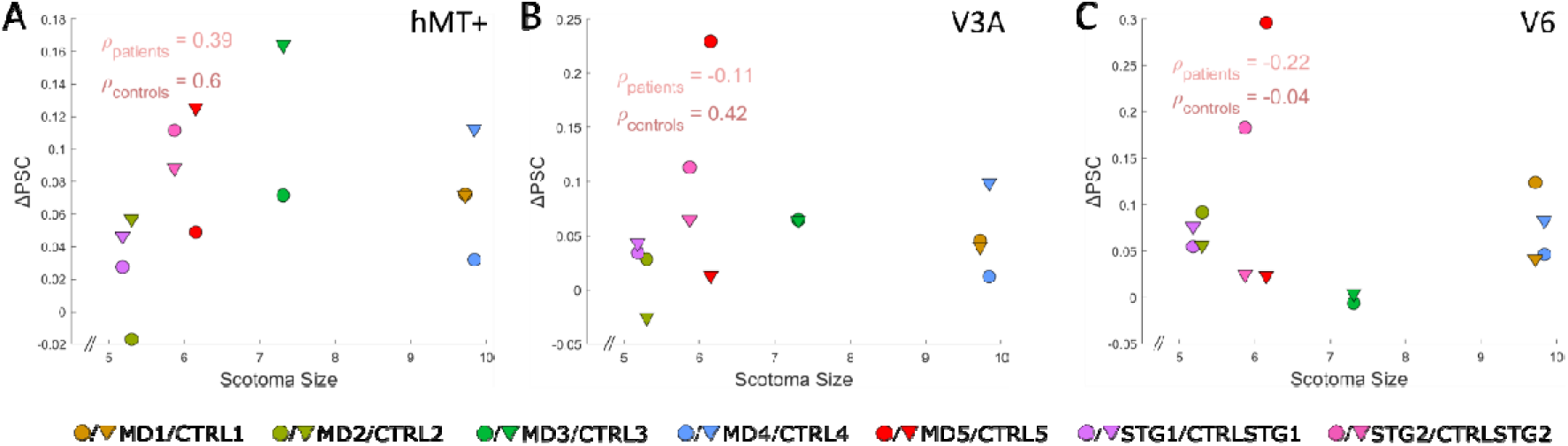
Differences between the average percentages of signal changes (PSCs) during the translational motion and control (static dots) conditions in the hMT+ (A), V3A (B) and V6 (C) ROIs as a function of the scotoma size, given by the radius of the associated disk (see the ‘*Methods*’ section). The corresponding Spearman correlation coefficients rho (⍴) are shown. None of these correlations were significant (p-values > 0.05). For patient MD5 (whose scotoma is elongated, see supplementary Figure 1) and his control, we used the semi minor axis of the associated ellipse for the scotoma size in this Figure. We obtained similar results when using the semi major axis of the ellipse.

---

**Supplementary Table 2-1:** Talairach coordinates of pSTS observed in patients and controls, the calculated mean for both patients and controls, and the coordinates reported in Barton and Brewer (2017). The ‘*’ symbol indicates the coordinates for which a more permissive threshold has been used, i.e p<10-2.

|  |  | pSTS |
| --- | --- | --- |
| MD1 | LH | X |
|  | RH | X |
| MD2 | LH | X |
|  | RH | 51; -48; 10* |
| MD3 | LH | -47; -46; 2 |
|  | RH | 53; -51; 5 |
| MD4 | LH | -47; -47; 9 |
|  | RH | 56; -40; 5 |
| MD5 | LH | X |
|  | RH | X |
| STG1 | LH | -39; -49; 11* |
|  | RH | 51; -43; 12* |
| STG2 | LH | -49; -50; 5 |
|  | RH | 54; -41; 6 |
| CTRL1 | LH | X |
|  | RH | X |
| CTRL2 | LH | -51; -40; 3 |
|  | RH | 52; -33; 5 |
| CTRL3 | LH | -46; -38; -1 |
|  | RH | 43; -42; -1 |
| CTRL4 | LH | X |
|  | RH | 55; -35; 10 |
| CTRL5 | LH | X |
|  | RH | 44; -35; 1 |
| CTRLSTG1 | LH | -45; -49; 10 |
|  | RH | 53; -37; 11 |
| CTRLSTG2 | LH | -47; -48; 5 |
|  | RH | 40; -54; 12 |
| Average patients | LH | -46±4; -48±8; 7±4 |
|  | RH | 53±3; -45±3; 8±2 |
| Average controls | LH | -47±2; -44±1; 4±2 |
|  | RH | 48±1; -39±1; 6±1 |
| Literature | LH | -56; -47; 8 |
|  | RH | 52; -49; 15 |

## References

Aedo-Jury, F., Cottereau, B. R., Celebrini, S., & Séverac Cauquil, A. (2020). Antero-Posterior vs. Lateral Vestibular Input Processing in Human Visual Cortex. Frontiers in Integrative Neuroscience, 14, 43. 10.3389/fnint.2020.00043

Audurier, P., Héjja-Brichard, Y., De Castro, V., Kohler, P. J., Norcia, A. M., Durand, J.-B., & Cottereau, B. R. (2022). Symmetry Processing in the Macaque Visual Cortex. Cerebral Cortex, 32(10), 2277–2290. 10.1093/cercor/bhab358

Bach, M. (2007). The Freiburg Visual Acuity Test-variability unchanged by post-hoc re-analysis. Graefe’s Archive for Clinical and Experimental Ophthalmology = Albrecht Von Graefes Archiv Fur Klinische Und Experimentelle Ophthalmologie, 245(7), 965–971. 10.1007/s00417-006-0474-4

Baker, C. I. (2005). Reorganization of Visual Processing in Macular Degeneration. Journal of Neuroscience, 25(3), 614–618. 10.1523/JNEUROSCI.3476-04.2005

Barton, B., & Brewer, A. A. (2017). Visual Field Map Clusters in High-Order Visual Processing: Organization of V3A/V3B and a New Cloverleaf Cluster in the Posterior Superior Temporal Sulcus. Frontiers in Integrative Neuroscience, 11. 10.3389/fnint.2017.00004

Benson, N. C., & Winawer, J. (2018). Bayesian analysis of retinotopic maps. eLife, 7, e40224. 10.7554/eLife.40224

Bremmer, F., Schlack, A., Shah, N. J., Zafiris, O., Kubischik, M., Hoffmann, K.-P., Zilles, K., & Fink, G. R. (2001). Polymodal Motion Processing in Posterior Parietal and Premotor Cortex. Neuron, 29(1), 287–296. 10.1016/S0896-6273(01)00198-2

Burnat, K., Hu, T.-T., Kossut, M., Eysel, U. T., & Arckens, L. (2017). Plasticity Beyond V1: Reinforcement of Motion Perception upon Binocular Central Retinal Lesions in Adulthood. Journal of Neuroscience, 37(37), 8989–8999. 10.1523/JNEUROSCI.1231-17.2017

Calford, M. B., Chino, Y. M., Das, A., Eysel, U. T., Gilbert, C. D., Heinen, S. J., Kaas, J. H., & Ullman, S. (2005). Rewiring the adult brain. Nature, 438(7065), E31ZE3. 10.1038/nature04359

Carvalho, J., Invernizzi, A., Martins, J., Renken, R. J., & Cornelissen, F. W. (2022). Local neuroplasticity in adult glaucomatous visual cortex. Scientific Reports, 12(1), 21981. 10.1038/s41598-022-24709-1

Chakraborty, A., Tran, T. T., Silva, A. E., Giaschi, D., & Thompson, B. (2021). Continuous theta burst TMS of area MT+ impairs attentive motion tracking. The European Journal of Neuroscience, 54(9), 7289–7300. 10.1111/ejn.15480

Christen, M., & Abegg, M. (2017). The effect of magnification and contrast on reading performance in different types of simulated low vision. Journal of Eye Movement Research, 10(2). 10.16910/jemr.10.2.5

Contemori, G., Maniglia, M., Guénot, J., Soler, V., Cherubini, M., Cottereau, B. R., & Trotter, Y. (2024). tRNS boosts visual perceptual learning in participants with bilateral macular degeneration. Frontiers in Aging Neuroscience, 16, 1326435. 10.3389/fnagi.2024.1326435

Dilks, D. D., Baker, C. I., Peli, E., & Kanwisher, N. (2009). Reorganization of visual processing in macular degeneration is not specific to the « preferred retinal locus ». The Journal of Neuroscience: The Official Journal of the Society for Neuroscience, 29(9), 2768–2773. 10.1523/JNEUROSCI.5258-08.2009

Dumoulin, S. O. (2000). A New Anatomical Landmark for Reliable Identification of Human Area V5/MT: A Quantitative Analysis of Sulcal Patterning. Cerebral Cortex, 10(5), 454–463. 10.1093/cercor/10.5.454

Dumoulin, S. O., Baker, C. L., Jr, Hess, R. F., & Evans, A. C. (2003). Cortical Specialization for Processing First-and Second-order Motion. Cerebral Cortex, 13(12), 1375–1385. 10.1093/cercor/bhg085

Esteban, O., Birman, D., Schaer, M., Koyejo, O. O., Poldrack, R. A., & Gorgolewski, K. J. (2017). MRIQC-: Advancing the automatic prediction of image quality in MRI from unseen sites. PLOS ONE, 12(9), e0184661. 10.1371/journal.pone.0184661

Fleming, L. L., Defenderfer, M. K., Demirayak, P., Stewart, P., Decarlo, D. K., & Visscher, K. M. (2024). Impact of Deprivation and Preferential Usage on Functional Connectivity Between Early Visual Cortex and Category-Selective Visual Regions. Human Brain Mapping, 45(17), e70064. 10.1002/hbm.70064

Fletcher, D. C., & Schuchard, R. A. (1997). Preferred Retinal Loci Relationship to Macular Scotomas in a Low-vision Population. Ophthalmology, 104(4), 632–638. 10.1016/S0161-6420(97)30260-7

Goebel, R., Esposito, F., & Formisano, E. (2006). Analysis of functional image analysis contest (FIAC) data with brainvoyager QX: From single-subject to cortically aligned group general linear model analysis and self-organizing group independent component analysis. Human Brain Mapping, 27(5), 392–401. 10.1002/hbm.20249

Graziano, M., Andersen, R., & Snowden, R. (1994). Tuning of MST neurons to spiral motions. The Journal of Neuroscience, 14(1), 54–67. 10.1523/JNEUROSCI.14-01-00054.1994

Grossman, E. D., & Blake, R. (2002). Brain Areas Active during Visual Perception of Biological Motion. Neuron, 35(6), 1167–1175. 10.1016/S0896-6273(02)00897-8

Guénot, J., Trotter, Y., Delaval, A., Baurès, R., Soler, V., & Cottereau, B. R. (2023). Processing of translational, radial and rotational optic flow in older adults. Scientific Reports, 13(1), 15312. 10.1038/s41598-023-42479-2

Guénot, J., Trotter, Y., Fricker, P., Cherubini, M., Soler, V., & Cottereau, B. R. (2022). Optic Flow Processing in Patients With Macular Degeneration. Investigative Ophthalmology & Visual Science, 63(12), 21. 10.1167/iovs.63.12.21

Gupta, A., Mesik, J., Engel, S. A., Smith, R., Schatza, M., Calabrèse, A., van Kuijk, F. J., Erdman, A. G., & Legge, G. E. (2018). Beneficial Effects of Spatial Remapping for Reading With Simulated Central Field Loss. Investigative Ophthalmology & Visual Science, 59(2), 1105–1112. 10.1167/iovs.16-21404

Hassan, S. E., Lovie-Kitchin, J. E., & Woods, A. R. L. (2002). Vision and Mobility Performance of Subjects with Age-Related Macular Degeneration: Optometry and Vision Science, 79(11), 697–707. 10.1097/00006324-200211000-00007

Hernowo, A. T., Prins, D., Baseler, H. A., Plank, T., Gouws, A. D., Hooymans, J. M. M., Morland, A. B., Greenlee, M. W., & Cornelissen, F. W. (2014). Morphometric analyses of the visual pathways in macular degeneration. Cortex, 56, 99–110. 10.1016/j.cortex.2013.01.003

Huk, A. C., Dougherty, R. F., & Heeger, D. J. (2002). Retinotopy and Functional Subdivision of Human Areas MT and MST. The Journal of Neuroscience, 22(16), 7195–7205. 10.1523/JNEUROSCI.22-16-07195.2002

Kolster, H., Peeters, R., & Orban, G. A. (2010). The Retinotopic Organization of the Human Middle Temporal Area MT/V5 and Its Cortical Neighbors. Journal of Neuroscience, 30(29), 9801–9820. 10.1523/JNEUROSCI.2069-10.2010

Kuyk, T., & Elliott, J. L. (1999). Visual factors and mobility in persons with age-related macular degeneration. Journal of Rehabilitation Research and Development, 36(4), 303–312.

Li, J. Q., Welchowski, T., Schmid, M., Mauschitz, M. M., Holz, F. G., & Finger, R. P. (2020). Prevalence and incidence of age-related macular degeneration in Europe: A systematic review and meta-analysis. The British Journal of Ophthalmology, 104(8), 1077–1084. 10.1136/bjophthalmol-2019-314422

Little, D., Thulborn, K., & Szlyk, J. (2008). An FMRI study of saccadic and smooth-pursuit eye movement control in patients with age-related macular degeneration. Investigative Ophthalmology & Visual Science, 49(4). 10.1167/iovs.07-0372

Maniglia, M., Soler, V., Cottereau, B., & Trotter, Y. (2018). Spontaneous and training-induced cortical plasticity in MD patients: Hints from lateral masking. Scientific Reports, 8(1), 90. 10.1038/s41598-017-18261-6

Marron, J. A., & Bailey, I. L. (1982). Visual Factors and Orientation-Mobility Performance: Optometry and Vision Science, 59(5), 413–426. 10.1097/00006324-198205000-00009

Masuda, Y., Dumoulin, S. O., Nakadomari, S., & Wandell, B. A. (2008). V1 projection zone signals in human macular degeneration depend on task, not stimulus. Cerebral Cortex (New York, N.Y.: 1991), 18(11), 2483-2493. 10.1093/cercor/bhm256

Masuda, Y., Takemura, H., Terao, M., Miyazaki, A., Ogawa, S., Horiguchi, H., Nakadomari, S., Matsumoto, K., Nakano, T., Wandell, B. A., & Amano, K. (2021). V1 Projection Zone Signals in Human Macular Degeneration Depend on Task Despite Absence of Visual Stimulus. Current Biology, 31(2), 406–412.e3. 10.1016/j.cub.2020.10.034

McKeefry, D. J., Burton, M. P., Vakrou, C., Barrett, B. T., & Morland, A. B. (2008). Induced deficits in speed perception by transcranial magnetic stimulation of human cortical areas V5/MT+ and V3A. The Journal of Neuroscience: The Official Journal of the Society for Neuroscience, 28(27), 6848–6857. 10.1523/JNEUROSCI.1287-08.2008

Morland, A. B. (2015). Organization of the Central Visual Pathways Following Field Defects Arising from Congenital, Inherited, and Acquired Eye Disease. Annual Review of Vision Science, 1(1), 329–350. 10.1146/annurev-vision-082114-035600

Musel, B., Hera, R., Chokron, S., Alleysson, D., Chiquet, C., Romanet, J.-P., Guyader, N., & Peyrin, C. (2011). Residual abilities in age-related macular degeneration to process spatial frequencies during natural scene categorization. Visual Neuroscience, 28(6), 529–541. 10.1017/S0952523811000435

Papanikolaou, A., Keliris, G. A., Papageorgiou, T. D., Shao, Y., Krapp, E., Papageorgiou, E., Stingl, K., Bruckmann, A., Schiefer, U., Logothetis, N. K., & Smirnakis, S. M. (2014). Population receptive field analysis of the primary visual cortex complements perimetry in patients with homonymous visual field defects. Proceedings of the National Academy of Sciences, 111(16). 10.1073/pnas.1317074111

Peyrin, C., Ramanoël, S., Roux-Sibilon, A., Chokron, S., & Hera, R. (2017). Scene perception in age-related macular degeneration: Effect of spatial frequencies and contrast in residual vision. Vision Research, 130, 36–47. 10.1016/j.visres.2016.11.004

Pigarev, I. N., & Rodionova, E. I. (1998). Two visual areas located in the middle suprasylvian gyrus (cytoarchitectonic field 7) of the cat’s cortex. Neuroscience, 85(3), 717–732. 10.1016/S0306-4522(97)00642-8

Pilly, P. K.,& Seitz, A. R. (2009). What a difference a parameter makes: A psychophysical comparison of random dot motion algorithms. Vision Research, 49(13), 15991Z–1612. 10.1016/j.visres.2009.03.019

Pitzalis, S., Galletti, C., Huang, R.-S., Patria, F., Committeri, G., Galati, G., Fattori, P., & Sereno, M. I. (2006). Wide-Field Retinotopy Defines Human Cortical Visual Area V6. The Journal of Neuroscience, 26(30), 7962–7973. 10.1523/JNEUROSCI.0178-06.2006

Pitzalis, S., Sereno, M. I., Committeri, G., Fattori, P., Galati, G., Patria, F., & Galletti, C. (2010). Human v6: The medial motion area. *Cerebral Cortex (New York*, N.Y.: 1991), *20*(2), 411-424. 10.1093/cercor/bhp112

Poldrack, R. A. (2007). Region of interest analysis for fMRI. Social Cognitive and Affective Neuroscience, 2(1), 67–70. 10.1093/scan/nsm006

Priebe, N. J., Lisberger, S. G., & Movshon, J. A. (2006). Tuning for Spatiotemporal Frequency and Speed in Directionally Selective Neurons of Macaque Striate Cortex. The Journal of Neuroscience, 26(11), 2941–2950. 10.1523/JNEUROSCI.3936-05.2006

Raftery, A. E. (1995). Bayesian Model Selection in Social Research. Sociological Methodology, 25, 111. 10.2307/271063

Ramanoël, S., Chokron, S., Hera, R., Kauffmann, L., Chiquet, C., Krainik, A., & Peyrin, C. (2018). Age-related macular degeneration changes the processing of visual scenes in the brain. Visual Neuroscience, 35, E006. 10.1017/S0952523817000372

Shao, Y., Keliris, G. A., Papanikolaou, A., Fischer, M. D., Zobor, D., Jägle, H., Logothetis, N. K., & Smirnakis, S. M. (2013). Visual cortex organisation in a macaque monkey with macular degeneration. European Journal of Neuroscience, 38(10), 3456–3464. 10.1111/ejn.12349

Silson, E. H., Baker, C. I., Aleman, T. S., Maguire, A. M., Bennett, J., & Ashtari, M. (2023). Motion-selective areas V5/MT and MST appear resistant to deterioration in choroideremia. NeuroImage: Clinical, 38, 103384. 10.1016/j.nicl.2023.103384

Strong, S. L., Silson, E. H., Gouws, A. D., Morland, A. B., & McKeefry, D. J. (2017). A Direct Demonstration of Functional Differences between Subdivisions of Human V5/MT+. Cerebral Cortex (New York, NY), 27(1), 1–10. 10.1093/cercor/bhw362

Szlyk, J. P., & Little, D. M. (2009). An FMRI study of word-level recognition and processing in patients with age-related macular degeneration. Investigative Ophthalmology & Visual Science, 50(9), 4487–4495. 10.1167/iovs.08-2258

Talairach, J., & Tournoux, P. (1988). Co-planar Stereotaxic Atlas of the Human Brain-: 3-dimensional Proportional System: an Approach to Cerebral Imaging. G. Thieme.

Tarita-Nistor, L., González, E. G., Markowitz, S. N., Lillakas, L., & Steinbach, M. J. (2008). Increased Role of Peripheral Vision in Self-Induced Motion in Patients with Age-Related Macular Degeneration. Investigative Ophthalmology & Visual Science, 49(7), 3253–3258. 10.1167/iovs.07-1290

Tootell, R. B. H., Mendola, J. D., Hadjikhani, N. K., Ledden, P. J., Liu, Arthur. K., Reppas, J. B., Sereno, M. I., & Dale, A. M. (1997). Functional Analysis of V3A and Related Areas in Human Visual Cortex. The Journal of Neuroscience, 17(18), 7060–7078. 10.1523/JNEUROSCI.17-18-07060.1997

Tootell, R. B. H., Reppas, J. B., Dale, A. M., Look, R. B., Sereno, M. I., Malach, R., Brady, T. J., & Rosen, B. R. (1995). Visual motion aftereffect in human cortical area MT revealed by functional magnetic resonance imaging. Nature, 375(6527), 139–141. 10.1038/375139a0

Tootell, R., Reppas, J., Kwong, K., Malach, R., Born, R., Brady, T., Rosen, B., & Belliveau, J. (1995). Functional analysis of human MT and related visual cortical areas using magnetic resonance imaging. The Journal of Neuroscience, 15(4), 3215–3230. 10.1523/JNEUROSCI.15-04-03215.1995

Tran, T. H. C., Guyader, N., Guerin, A., Despretz, P., & Boucart, M. (2011). Figure Ground Discrimination in Age-Related Macular Degeneration. Investigative Opthalmology & Visual Science, 52(3), 1655. 10.1167/iovs.10-6003

Villeneuve, M. Y., Ptito, M., & Casanova, C. (2006). Global motion integration in the postero-medial part of the lateral suprasylvian cortex in the cat. Experimental Brain Research, 172(4), 485–497. 10.1007/s00221-006-0357-2

Watson, A. B., & Pelli, D. G. (1983). Quest: A Bayesian adaptive psychometric method. Perception & Psychophysics, 33(2), 113–120. 10.3758/BF03202828

Westeneng-van Haaften, S. C., Boon, C. J. F., Cremers, F. P. M., Hoefsloot, L. H., Den Hollander, A. I., & Hoyng, C. B. (2012). Clinical and Genetic Characteristics of Late-onset Stargardt’s Disease. Ophthalmology, 119(6), 1199–1210. 10.1016/j.ophtha.2012.01.005

Wood, J. M., Lacherez, P. F., Black, A. A., Cole, M. H., Boon, M. Y., & Kerr, G. K. (2009). Postural Stability and Gait among Older Adults with Age-Related Maculopathy. Investigative Opthalmology & Visual Science, 50(1), 482. 10.1167/iovs.08-1942

Zeki, S., Watson, J. D., Lueck, C. J., Friston, K. J., Kennard, C., & Frackowiak, R. S. (1991). A direct demonstration of functional specialization in human visual cortex. The Journal of Neuroscience: The Official Journal of the Society for Neuroscience, 11(3), 641–649. 10.1523/JNEUROSCI.11-03-00641.1991

